# Ultraplex microscopy: versatile highly-multiplexed molecular labeling and imaging across scale and resolution

**DOI:** 10.1101/2024.08.17.605585

**Authors:** Janeth Pérez-Garza, Jairo Orea, Zachary Deane, Gianna Raimondi, Rebecca Tripp, Imani Charles, Linnaea Ostroff

## Abstract

The molecular organization of cells and tissue is challenging to study due to the inefficiency of multiplexed molecular labeling methods and the limited options for combining microscopy modalities in a single specimen, especially when high spatial resolution is needed. Here we describe ultraplex microscopy, which combines serial multiplexing, ultrathin sectioning, and reversible embedding to circumvent incompatibilities between labeling and imaging techniques, enhance resolution, and expand multiplexing capacity within and across modalities. Samples can be labeled with antibodies, RNA probes, and tissue stains for imaging by brightfield, epifluorescence, super-resolution, and electron microscopy without specialized reagents or materials. We demonstrate applications in brain tissue including molecular profiling of single cells and axonal boutons, high-resolution molecular colocalization, and correlative imaging of fluorescent proteins with confocal and ultraplex microscopy. The power and versatility of ultraplex microscopy will be valuable in addressing currently intractable experimental questions in many systems and contexts.

## Introduction

Brain cells are morphologically and molecularly heterogeneous^1–3^, and their complex patterns of gene expression and protein localization are dynamically regulated on a subcellular scale^3^. Understanding the brain’s anatomical organization – as well as how it varies with species, age, sex, experience, and disease – will require mapping tissue structure alongside large numbers of molecules at high resolution. Routine methods are available for localizing various molecules at scales from millimeters to microns and for imaging tissue with nanometer resolution, but it is currently impossible to combine a full complement of molecular labeling and imaging methods on a single sample. Some constraints on multiplexed molecular labeling can be mitigated; cyclic labeling strategies ^4,5^ increase the number of imaging channels available for immunofluorescence (IF) or fluorescence *in situ* hybridization (FISH) and underlie some spatial transcriptomics techniques^6^, while cross-reactivity between antibodies in IF can be avoided by replacing standard reagents with customized quenchable^7,8^ or releasable tags^9,10^. These approaches do not address incompatibility between labeling and imaging methods, however. Antibody labeling procedures can degrade RNA while FISH protocols destroy protein antigens, such that some protein and RNA targets cannot be labeled in the same sample^11^. Because the correlation between gene transcripts and proteins is low and varies between brain regions^12,13^, this is a serious problem. Preservation of tissue morphology is often at odds with molecular labeling, as tissue must be permeabilized or digested to allow probes access to their targets. Electron microscopy (EM) cannot be combined with RNA labeling for this reason, and combining EM with antibody labeling often requires compromising the results of both.^14,15^

A potentially powerful strategy for avoiding protocol incompatibility is serial multiplexing, in which serial sections of a sample are used for parallel, independent experiments. In theory, arbitrary combinations of molecular labels, staining methods, and imaging modalities can be applied to single cells or even organelles by imaging them across differently stained sections. Serial multiplexing has a major practical barrier: for the strategy to be effective and efficient, sections must be substantially thinner than the objects of interest yet robust enough for handling and reconstruction. Vibratome sections of fixed brain tissue ≥ 40 µm thick have sufficient mechanical integrity for single cells to be followed from one section to the next ^16–18^, but thinner sections and cryosections are subject to too much distortion^16^. Neurons are smaller than 40 µm in diameter and thus can appear on no more than two sections, so the benefits of serial multiplexing are minimal.

Samples embedded in plastic resin for EM can be cut into hundreds or even thousands of sturdy ultrathin (∼50 nm) sections for volume reconstruction of cells and organelles^19–21^, and serial multiplexing has been used at the EM level to identify neurons containing amino acid neurotransmitters^22–24^ and peptide hormones^25^. Outside of these targets^26^, however, most molecules are difficult to detect on conventional EM sections, in large part due to the presence of embedding resin. Antigenicity can be improved by chemical etching or the use of alternative resins^27–31^, and EM resin sections have been used for immunolabeling at the light microscopy level^28^ in cyclic^32–34^ and serial multiplexed^29^ designs that exploit the enhanced spatial resolution they offer^33,34^. Sensitivity is still low, however, as evidenced by the use of thicker sections^29,35^, high concentrations of antibodies^32,33^, and specialized screening for antibodies compatible with resin sections^36^.

Serial multiplexing across ultrathin sections could allow otherwise impossible combinations of labeling and imaging modalities to be applied to a single sample, and therefore has the potential to reveal molecular localization in depth and detail that are currently unobtainable. Ultrathin sections must be compatible with molecular labeling for this approach to work, however, meaning that samples cannot be embedded in EM resin. Paraffin wax embedding is routinely used to prepare tissue sections for light microscopy, and in contrast to EM resins, paraffin sections are a standard format for IF and FISH^37,38^. A key difference between paraffin wax and EM resin is that paraffin wax is removed from sections before labeling, so we developed a reversible embedding method to allow ultrathin sections to be labeled in the absence of an embedding medium. Here we demonstrate that the combination of reversible embedding, ultrathin sectioning, and serial multiplexing, which we term ultraplex microscopy, preserves compatibility with a range of molecular labeling and imaging methods. Further, we demonstrate its use in applications ranging from profiling of single cells and axonal boutons to correlative confocal imaging of fluorescent proteins.

## Results

### Reversible embedding for tissue preservation and ultrathin sectioning

The goal for the ultraplex workflow was to stabilize tissue for serial ultrathin sectioning while preserving its compatibility with a range of molecular labeling, staining, and imaging methods (Figure 1A). An ideal embedding medium would be a hard plastic that has excellent ultrathin sectioning properties, is soluble for gentle removal, has good optical clarity, and does not interact with tissue molecules. Polymethacrylates are highly customizable due to the variety of available monomers^39,40^ and are commonly used for embedding, both in EM resins^31^ and removable histology media^41–44^. We optimized a polymethacrylate formula to produce high-quality ultrathin sections that could be deplasticized with acetone, and tested its performance by embedding samples of fixed amygdala from adult rats.

**Figure 1.**
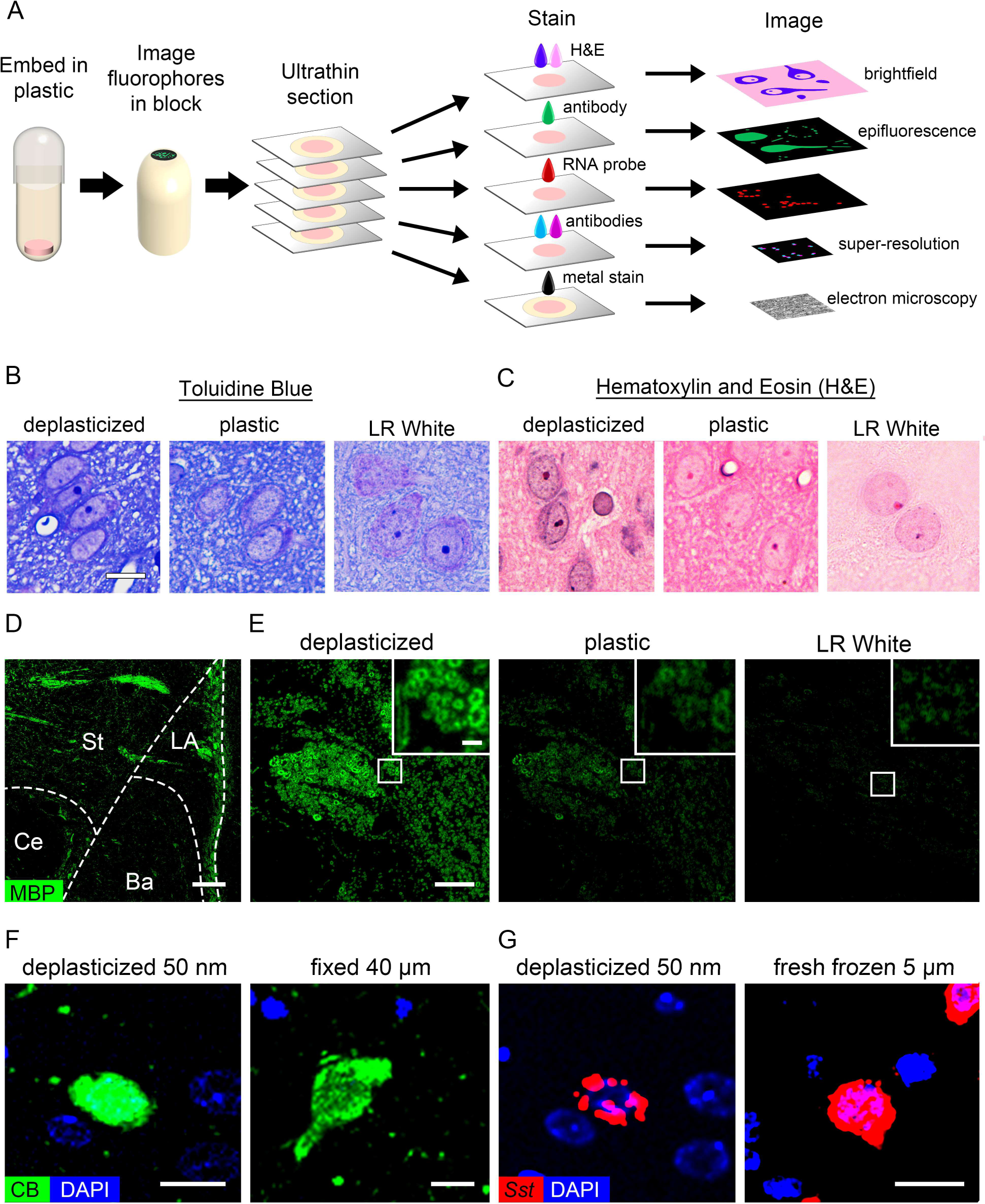
Ultraplex microscopy implementation with reversible embedding. A) Schematic of ultraplex workflow. B) Thick (1 µm) sections of rat amygdala in deplasticized (left), plastic (center) and LR White (right) stained with toluidine blue. C) Hematoxylin and eosin staining (H&E) of 2 µm sections of rat amygdala in deplasticized (left), plastic (center) and LR White (right). D-E) Immunofluorescence for myelin basic protein on 50 nm sections of rat amygdala. D) A deplasticized section imaged at 10X. E) The boxed region in (D) imaged at 60X (left), the same region of an adjacent section labeled with the plastic intact (center), and a similar region in an LR White section (right). Insets show enlargements of the boxed regions. F) Immunofluorescence for calbindin on a 50 nm deplasticized section (left) and a 40 µm vibratome section of rat amygdala (right). G) FISH labeling with a somatostatin probe on a 50 nm deplasticized section (left) and a 5 µm cryosection of fresh-frozen tissue (right). In (F) and (G) deplasticized sections were imaged in a widefield microscope at 20X and single 0.5 µm optical sections of vibratome and confocal sections were imaged at 40X in a confocal microscope. LA: lateral amygdala; Ba: Basal amygdala; Ce: Central amygdala; St: Striatum; MBP: myelin basic protein; CB: calbindin; Sst: somatostatin mRNA. Scale bar = 10 µm in (B-C), 100 µm in (D), 20 µm in (E), 2 µm in (E, inset), and 10 µm in (F-G).

We first examined preservation of tissue structure using histological stains, starting with toluidine blue, which is commonly used to assess gross morphology in thick EM resin sections by light microscopy. On 1 μm sections, toluidine blue staining was denser on deplasticized sections than on sections with intact plastic or sections of LR White, an EM resin often recommended for immunolabeling^31^ (Figure 1B). Tissue features were also more difficult to discern at low magnification on deplasticized sections (Figure S1A). Whereas toluidine blue staining is optimized for producing contrast on EM resin sections, hematoxylin and eosin (H&E) staining is standard for visualizing tissue structure on dewaxed paraffin sections. H&E produced good contrast on deplasticized sections, but very little hematoxylin staining appeared on plastic sections and neither component of the stain was taken up effectively by LR White sections (Figure 1C, Figure S1B).

Our embedding formula sectioned easily at 50 nm and formed ordered ribbons that facilitated collecting serial sections in order, unlike LR White (Figure S2). It is possible that ultrathin sections could disintegrate or detach from the substrate during labeling if they are not stabilized by plastic, so we tested IF on deplasticized 50 nm sections compared to sections with intact plastic and LR White sections using the same antibody concentration, labeling conditions and imaging settings (detailed settings for this and subsequent experiments are given in Table S1). An antibody to myelin basic protein revealed the expected pattern of white matter at low magnification (Figure 1D), confirming that the tissue was robust and could produce strong signal despite its thinness, and at 60X rings of myelin were easily visible on deplasticized sections but signal was faint on plastic and resin sections (Figure 1E).

### Detection of molecular cell-type markers on 50 nm sections

To establish whether labeling for endogenous molecular cell-type markers could be used to profile single cells on deplasticized 50 nm sections, we tested common interneuron markers in the rat amygdala. IF signal for calbindin filled cell bodies similarly on 50 nm sections and conventional 40 μm free-floating vibratome sections (Figure 1F). The same distribution of calbindin labeling in cells and processes was seen in both preparations at lower magnification (Figure S3A-B), and an antibody to parvalbumin produced similar results (Figure S3C-D). IF for calbindin was also stronger on deplasticized sections than plastic or LR White sections (Figure S3E-F), and no signal appeared when the primary antibody was omitted (Figure S3G-H).

Neuropeptides concentrate in axons, and because detection of peptidergic neurons with antibodies thus requires blocking axonal transport with colchicine^45^ we used FISH to detect these cells instead. A probe for somatostatin mRNA produced signal concentrated around nuclei on 50 nm sections, and a similar but somewhat more dense pattern was seen in cells on standard 5 μm cryosections (Figure 1G). The distribution of signal throughout the amygdala was the same on ultrathin and conventional sections, and probes for *Gad1* and parvalbumin mRNA also produced similar patterns, again with slightly lower density in cell bodies consistent with the lower section thickness (Figure S4A-F). FISH signal was weaker, although still specific, on plastic sections, while LR White tended to accumulate chunks of nonspecific signal even with a negative control probe (Figure S4G-J).

Glutaraldehyde was included in the primary fixative because it was necessary for immunolabeling GABA on 50 nm sections (Figure 2A), and in pilot tests we did not find that it impaired labeling of other targets. In contrast, glutaraldehyde had to be omitted from conventional preparations as it impaired IF and abolished FISH labeling. Glutaraldehyde also prevented the small wrinkles that sometimes formed during heat-induced antigen retrieval in samples fixed only with paraformaldehyde. Another advantage of deplasticized sections was that they are robust to antigen retrieval, which could not be used on plastic or LR White sections because it caused them to detach from the glass substrate.

**Figure 2.**
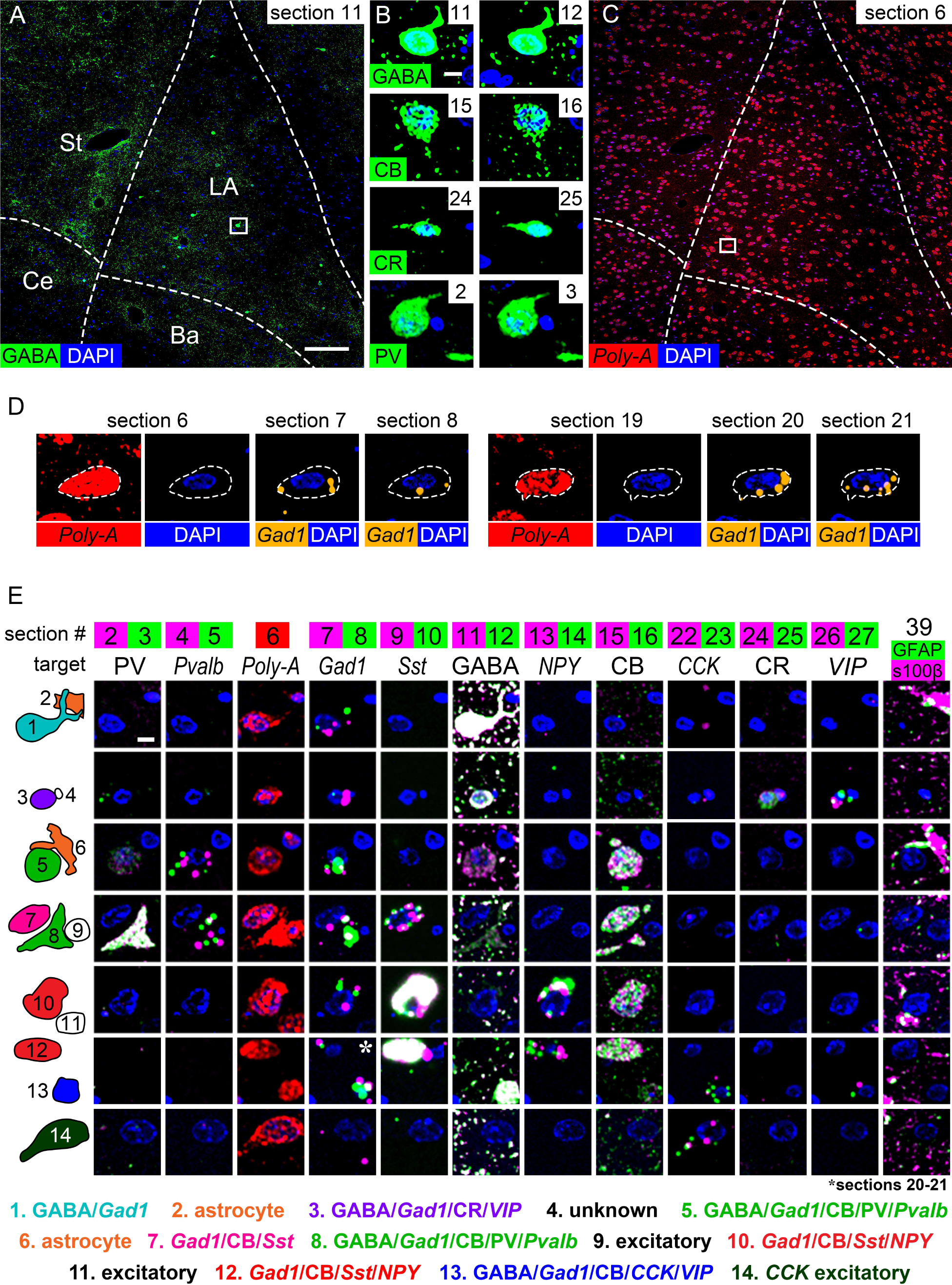
Serial multiplexed labeling of cell-type markers on resinless 50 nm sections. A) IF labeling for GABA on section #11 of 40. B) Single cells on pairs of adjacent sections labeled with the same antibody. C) FISH labeling with a poly-A mRNA probe on section #6 of 40. D) A single cell on six different sections, with the poly-A mRNA probe confirming that puncta of Gad1 mRNA fall within the cell’s region of mRNA-rich cytoplasm (dotted line). E) Examples of cells (rows) across selected sections (columns) labeled with various antibodies and mRNA probes. Labeling on pairs of sections is overlaid in each panel, except for the poly-A mRNA probe, which was applied to single sections, and IF for GFAP and s100β, which were labeled on the same sections. The cells in each row are numbered in the left column and labeling results are listed below. Labels in standard type indicate antibody targets and italics indicate target mRNA transcripts. All images in the figure were taken at 20X. PV: parvalbumin; CB: calbindin; CR: calretinin. Scale bar = 100 µm in (A,C), 5 µm in (B,D) and (E).

### Serial multiplexed molecular profiling at the single-cell level

Inhibitory interneurons in the rat basolateral amygdala complex are a useful test case for establishing an ultraplex workflow for single-cell profiling of molecular cell-type markers. Co-expression patterns of calcium-binding proteins and neuropeptides have been studied in these cells using double- and triple-label fluorescence and the mirror technique^46^, and we sought to detect some of the reported combinations across serial ultrathin sections of a single sample. Because most of the historic literature focuses on male subjects, we used males to facilitate comparison. A set of 40 serial 50 nm sections were collected on glass coverslips and labeled with antibodies or mRNA probes for several common markers (Figure S5). To facilitate labeling sections in parallel, a single IF protocol was used for all antibodies and most FISH labeling was performed with RNAscope, a commercial system whose probe design allows the same protocol to be used for all transcripts^47^. Imaging at 20X allowed for efficient acquisition of the entire lateral and basal amygdala while leaving single cells easily visible (Figure 2A-B).

To confirm consistent labeling across sections, all probes and antibodies were applied to pairs of adjacent sections within the series (Figure S5). Antibodies to GABA and the calcium binding proteins filled cell bodies (Figure 1F) and signal was reliably continuous across section pairs (Figure 2B). FISH signal was usually punctate and appeared in a peri-nuclear pattern, and signal density varied between probes and sometimes between cells (Figure S4). For probes with fewer puncta per cell, comparing adjacent sections was not always sufficient to resolve ambiguity because puncta did not always overlap with nuclear staining. To address this problem, some sections were labeled with a probe for poly-A mRNA (Figure 2C). The poly-A probe delineated a region of mRNA-rich cytoplasm surrounding each nucleus that was used to determine whether signal for other probes on nearby sections was localized within a given cell body (Figure 2D). When FISH probes were tested on sections varying from 50 nm to 2 µm in thickness, FISH signal density was predictably higher on thicker sections but labeling was present in the same cells across section thicknesses (Figure S6). Nevertheless, to increase sampling each FISH probe was repeated on two separate pairs of sections within the series (Figure 2D and Figure S5).

Examination of individual cells across serial sections (Figure 2E) revealed patterns of marker overlap consistent with previous reports^46^. Calbindin-positive cells were labeled for parvalbumin (Figure 2E, cells 5 and 8) or somatostatin (cells 7, 10, and 12), but not both. All somatostatin cells were labeled for calbindin, and some were labeled for *NPY* (cells 10 and 12) while others were not (cell 7). *VIP* labeling occurred with calretinin (cell 3) or *CCK* and calbindin (cell 13) in small cells with *Gad1* and GABA, and *CCK* also appeared in large cells without *Gad1* or GABA (cell 14). All cells labeled for GABA also contained *Gad1* mRNA, while cells with *Gad1* mRNA varied in the density of GABA labeling, from undetectable in cells containing somatostatin mRNA (7,10, and 12) to moderate in parvalbumin cells (5 and 8) to dense in calretinin/*VIP* cells (3), calbindin/*CCK*/*VIP* cells (13), and cells without any other markers (cell 1).

### Preservation of fluorescent proteins in ultrathin sections

Genetically-encoded fluorescent reporter proteins are often used to identify specific cells and axonal projections in brain tissue, and it is possible to preserve the fluorescence of reporter proteins in acrylic EM resins^48,49^. To test whether native fluorescence could be preserved for ultraplex microscopy, viral vectors were used to express GFP in the auditory cortex and tdTomato in the auditory thalamus of adult rats. The injected regions project to each other and to the lateral amygdala (Figure 3A), allowing for evaluation of fluorescence in the somatodendritic compartment as well as multiple long-range axonal projections. On deplasticized 50 nm sections of the cortical injection site, native fluorescence of GFP in cell bodies and tdTomato in the neuropil was readily visible at 10X and at higher magnification GFP could be seen filling larger processes suggestive of dendrites while tdTomato was in small processes consistent with axons (Figure 3B). Likewise, in the thalamic injection site tdTomato was visible in cell bodies and GFP was seen in small processes (Figure 3C).

**Figure 3.**
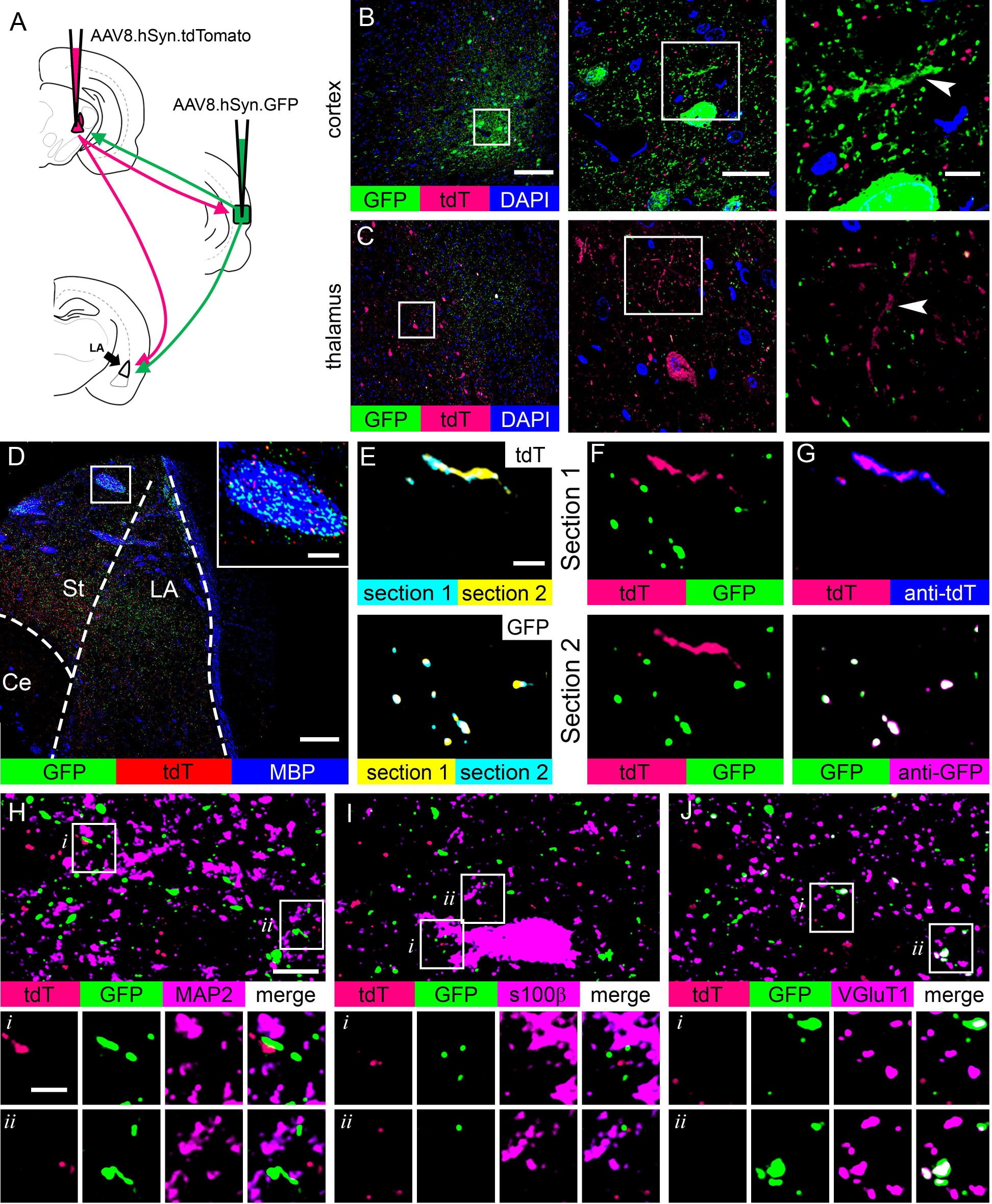
Preservation of native fluorescence in ultrathin sections. AAV vectors were used to express GFP and tdTomato in regions of the auditory cortex and auditory thalamus, respectively, that send projections to each other and to the lateral amygdala (LA). A) Schematic of the injections. B-C) Native fluorescence of GFP and tdTomato on 50 nm sections of the auditory cortex (B) and thalamus (C) from the same rat, imaged at 10X (left) and 60X (center). The boxed regions of the 60X images are enlarged (right) to show local dendrites (arrows) intermingled with axons from the distal injection site. D) A 50 nm section of LA from an injected rat imaged at 10X, showing IF for myelin basic protein (MBP) with native fluorescence of GFP and tdTomato. Inset shows enlargement of the boxed region. E-G) Native signal for tdTomato and GFP on two adjacent 50 nm sections of amygdala, with an antibody to tdTomato on section 1 and an antibody to GFP on section 2. E) Overlay of native signal for tdTomato (top) and GFP (bottom) on the two sections. F) Overlay of the native signals on each section. G) Antibody labeling for tdTomato on section 1 and GFP on section 2. H-J) 50 nm sections of amygdala immunolabeled for MAP2 (H), s100β (I), or VGluT1 (J) and imaged at 100X. Enlargements of boxed areas are shown in lower panels. Abbreviations in (D) in Figure 1. Scale bar = 100 µm in (B and C left, and D), 20 µm in (B,C center), 5 µm in (B,C right, H-J top panels), 20 µm in (D, inset), 2 µm in (E-G, H-J insets).

Cortical axons containing GFP and thalamic axons containing tdTomato were visible on 50 nm sections of amygdala at 10X, and both fluorophores appeared within white matter (identified by IF for myelin basic protein) as well as in the neuropil (Figure 3D). At high magnification, native signal for GFP and tdTomato could be followed across sections (Figure 3E) and the two fluorophores did not overlap with each other (Figure 3F), but antibody labeling for GFP and tdTomato overlapped with native fluorescence signals (Figure 3G).

### High-resolution colocalization of reporters and IF

Ultrathin sections offer a resolution advantage in light microscopy^33,34^, and the visibility of fluorophore-expressing axons provides an opportunity to observe the spatial precision of 50 nm sections in widefield fluorescence. On single sections, native fluorescence signals did not overlap with immunolabeling for MAP2 (Figure 3H), which localizes to dendrites, or s100β (Figure 3I), which localizes to astrocytic processes. Native fluorescence and IF signals did sometimes appear in close proximity, consistent with intermingling of axons, dendrites, and astrocytic processes in the neuropil. Some overlap was seen between immunolabeling for vesicular glutamate transporter 1 (VGluT1) and GFP, but not tdTomato (Figure 3J). This is expected, as cortical projection neurons are known to primarily express VGluT1 while thalamic projection neurons express vesicular glutamate transporter 2 (VGluT2).

As a test of signal co-localization, antibodies to VGluT1 and VGlut2 were applied separately to adjacent 50 nm sections of amygdala containing GFP-expressing cortical axons and tdTomato-expressing thalamic axons. GFP, tdTomato, and IF signal for both antibodies were visible throughout the lateral amygdala at 20X, and at 100X punctate patterns of all four signals were distributed in the neuropil (Figure S7). As predicted, overlapping puncta – presumably presynaptic boutons – of GFP and VGluT1 and of tdTomato and VGluT2 were visible at 100X (Figure 4A). Quantitative analysis of signal colocalization (Figure 4B) found that 32% of GFP signal colocalized with VGluT1, but almost none overlapped with VGluT2. Likewise, 42% of tdTomato signal colocalized with VGluT2, but none colocalized with VGluT1. A similar proportion of VGluT2 (35%) was colocalized with tdTomato, whereas only 5% of VGluT1 colocalized with GFP. The low percentage of VGluT1 puncta expressing GFP is likely due to a combination of VGluT1 expression in local amygdala neurons and GFP expression in a relatively small fraction of amgydala-projecting cortex.

**Figure 4.**
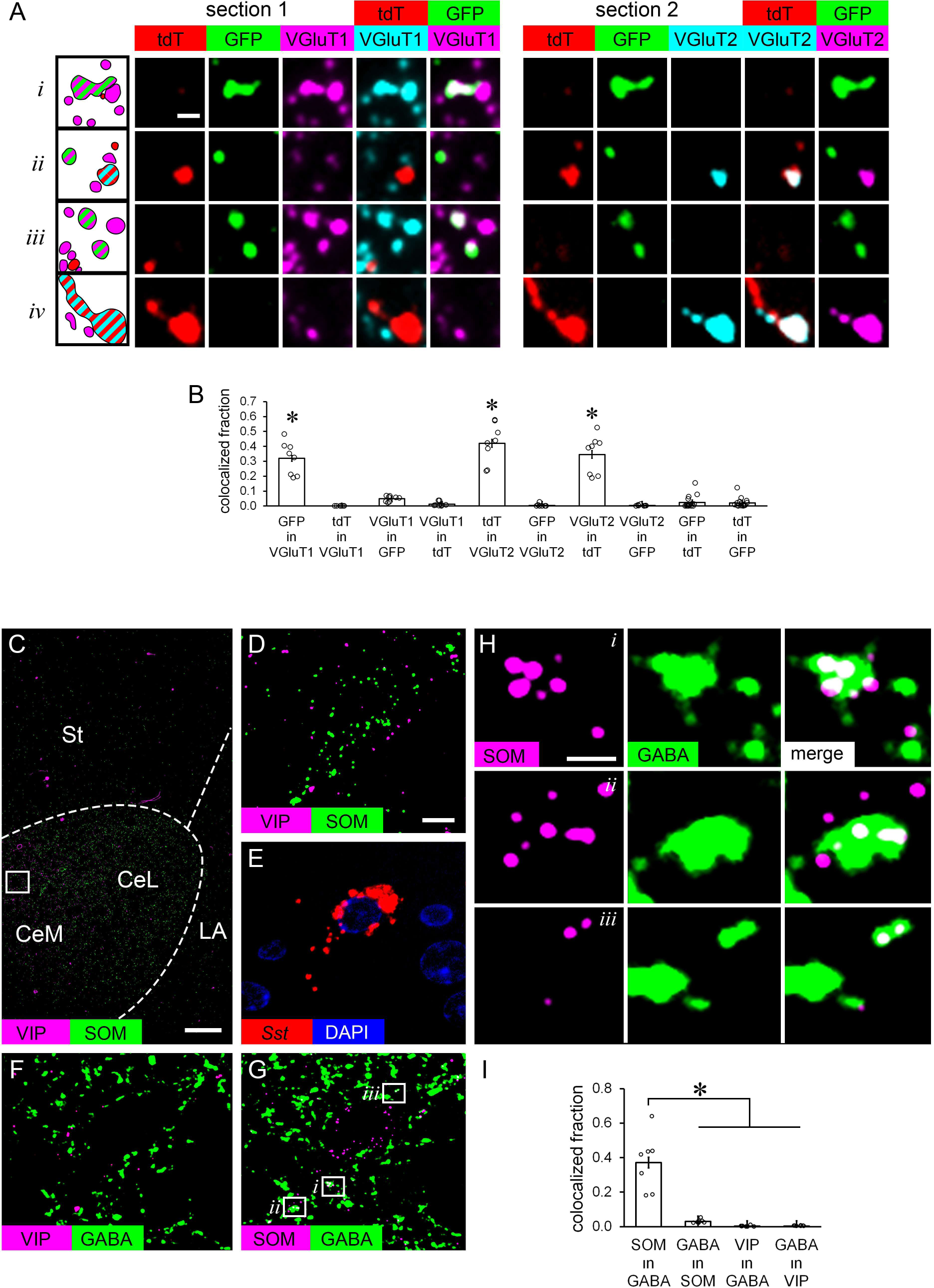
High-resolution co-localization of immunofluorescence in axons. A) Adjacent 50 nm sections of LA showing native fluorescence of tdTomato in axons from the auditory thalamus, GFP in axons from the auditory cortex, and immunofluorescence for both VGluT1 and VGluT2. B) Within-section co-localization analysis of tdTomato, GFP, and VGluT1 and VGluT2 immunofluorescence signals, given as the Manders coefficient representing the fraction of each signal colocalized with a second signal. An ANOVA found a main effect of signal fraction (F_(9,86)_ = 57.17, p < 0.001) and a Bonferroni post hoc test found that the fraction of GFP in VGlut1, tdT in VGluT2, and VGluT2 in tdT were each significantly different from the other signal pairs (* p < 0.001 in all cases) but not from each other. C) A 50 nm section of LA immunolabeled for VIP and somatostatin imaged at 20X. Note that VIP signal appears in blood vessels as well as neuropil. D) Boxed region of the section in (C) imaged at 100X. E-G) The region shown in (D) in three nearby serial sections, one labeled with a probe for somatostatin mRNA (E), one immunolabeled for VIP and GABA (F), and one immunolabeled for SOM and GABA (G). H) Enlargements of the boxed regions in (G). I) Within-section co-localization analysis of SOM and GABA immunofluorescence signals, as in (B). An ANOVA found a main effect of signal fraction (F_(3,24)_ = 34.70, p < 0.001) and a Bonferroni post hoc test found that the fraction of SOM in GABA was significantly different from all other signal pairs (* p < 0.001). LA: lateral amygdala, St: striatum, CeM: medial division of the central amygdala, CeL: lateral division of the central amygdala, VGluT1: vesicular glutamate transporter 1, VGluT2: vesicular glutamate transporter 2, VIP: vasoactive intestinal peptide, SOM: somatostatin, GABA: γ-aminobutyric acid. Error bars = s.e.m. Scale bar = 1 µm in (A), 100 µm in (C), 5 µm in (D-G), and 1 µm in (H).

### High-resolution colocalization of endogenous molecules

Having found that glutamate transporters can be localized with fluorophore-filled axons, we next asked whether endogenous molecules could be localized at a finer scale. Neuropeptides are a useful test case, as they are found in axons alongside small molecule neurotransmitters but in a different distribution; vesicles containing small molecule neurotransmitters form the main pools in presynaptic varicosities, while neuropeptides are found in large dense-core vesicles that are fewer in number and more widely distributed, occurring within synaptic vesicle pools, often at the periphery, and along the axon shaft^26,50^. The central nucleus of the amygdala contains dense axons expressing different neuropeptides; somatostatin is expressed by GABAergic neurons in the lateral division of the central amygdala (CeL) and somatostatin axons are also found in the medial division (CeM)^51^, while CeM also contains dense VIP fibers, the bulk of which enter from extrinsic sources via the medial forebrain bundle^52,53^ and do not appear to be GABAergic by electron microscopy^54^. Hence somatostatin and VIP should be intermingled in the neuropil, but only somatostatin should co-localize with GABA.

Double immunofluorescence labeling for both peptides on single sections revealed dense signal throughout the central nucleus that was visible even at 10X, with somatostatin and VIP concentrated in CeL and CeM, respectively, as expected (Figure 4C). At 100X, punctate signal for both peptides was seen throughout the neuropil at the boundary between CeL and CeM, and somatostatin puncta also filled some cell bodies (Figure 4D) that were confirmed to contain *Sst* mRNA by FISH labeling on nearby sections (Figure 4E). Unlike the punctate pattern of immunolabeling for the peptides, immunolabeling for GABA was widespread and diffuse (Figure 4F-G), with amorphous profiles of varying size presumably representing axons and dendrites.

Double immunolabeling on single sections revealed no apparent overlap between VIP and GABA (Figure 4F), but somatostatin sometimes overlapped with GABA (Figure 4G), with puncta visible within and adjacent to GABA profiles (Figure 4H). Quantitative analysis found that 37% of somatostatin signal was colocalized with GABA, but VIP labeling was not (Figure 4I). Only a small fraction of GABA labeling colocalized with somatostatin, consistent with the much higher absolute density of GABA signal throughout the neuropil. These results demonstrate the power of combining the labeling sensitivity of deplasticized sections with the imaging resolution provided by ultrathin sections.

### Cyclic immunofluorescence on resinless ultrathin sections

For molecular colocalization at high resolution or in very small structures, even ultrathin sections may be too thick for effective serial multiplexing and cyclic labeling may be necessary. Cyclic IF is challenging on ultrathin EM resin sections because antibody stripping treatments may detach sections from their substrates^32,55^, but we had noticed that our deplasticized sections remained on coverslips during antigen retrieval treatments while sections with plastic or LR White resin did not. To ensure that our sections could be used for cyclic IF, single 50 nm sections were double-labeled with primary antibodies to VGluT1 and VGlut2 and secondary antibodies conjugated to AlexaFluor-488 and AlexaFluor-647, then mounted and imaged as usual. Coverslips were removed and antibodies were stripped for re-staining using a protocol modified from Holderith and colleagues^32^. To assess whether labeling quality was maintained in the second round and whether antibodies had been fully removed, the fluorophores on the secondary antibodies were reversed in the second round, such that VGluT1 was detected with Alexa647 and VGluT2 with Alexa488 (Figure 5A). As a further test of antibody elution, the primary antibodies were omitted in the second round and the same secondary antibodies were applied. No signal was seen after the second labeling round(Figure 5B). Likewise, there was no signal overlap between rounds when the fluorophores were reversed (Figure 5C). When the average pixel intensity in the neuropil was compared across rounds and conditions, there was no difference between the first and second round when the same primaries were used, but signal was nearly abolished when the primaries were omitted (Figure 5D). Pixel intensity was significantly correlated between the two labeling rounds when the secondaries were reversed, but not when the primaries were omitted, and there was no correlation between imaging channels within rounds (Figure 5E).

**Figure 5.**
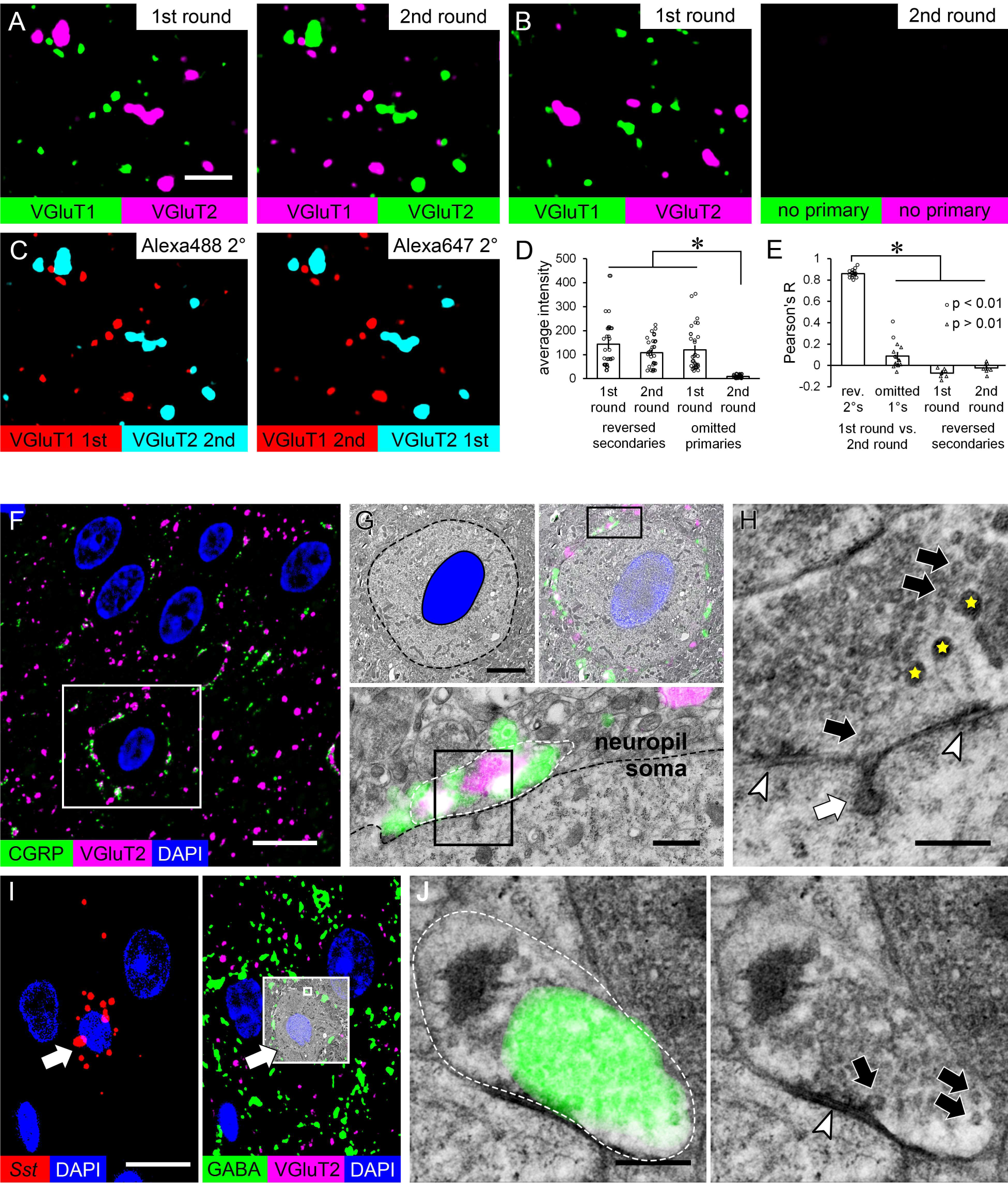
Cyclic immunofluorescence and serial multiplexed light and electron microscopy. A) Two rounds of immunofluorescence double-labeling on 50 nm sections of rat LA. In the first round, antibodies to VGluT1 and VGluT2 were detected with secondary antibodies conjugated to Alexa488 and Alexa647, respectively. In the second round, the same primary antibodies were applied but the fluorophores on the secondary antibodies were reversed. B) The same experiment, except that in the second round the primary antibodies were omitted and the same secondaries were applied in both rounds. C) The images in (A) with both rounds of each secondary overlaid. D) Comparison of average pixel intensity (out of a maximum of 4095 in the 12-bit images) of 50 µm^2^ ROIs in two labeling rounds, with the second round either with reversed secondary labels or omitted primaries. A two-way ANOVA found a significant interaction between labeling round and condition (F_(1,124)_ = 8.16, p = 0.005), and a Bonferroni post hoc test showed lower labeling between the second round with primaries omitted versus all other conditions (p < 0.001 in all cases). E) Correlation coefficients of pixel intensity in 50 µm^2^ ROIs. Comparisons were made of the same target channel between the two labeling rounds and between the two channels within each round in the reversed-secondaries condition. An ANOVA found that correlation coefficients between rounds were higher when the secondaries were reversed than when the primaries were omitted (F_(1,30)_ = 596.97, p < 0.0001). F-H) Two adjacent 50 nm sections of central amygdala, one immunolabeled and imaged by widefield fluorescence (F) and one imaged by TEM (G-H). F) Immunofluorescence for CGRP and VGluT2 imaged at 100X. G) The boxed region in (F) imaged by TEM at 2500X (top panels) and 10,000X (bottom panel). The top panels show the same image with the cell body indicated by a dashed line and the nucleus filled in blue (left) or with the IF overlaid (right). The bottom panel shows the boxed region in the top right panel. The dashed black line shows the boundary of the soma and the white dashed line encloses an axonal bouton. H) Enlarged view of the boxed region in the bottom panel of (G), showing large dense-core vesicles (yellow stars) and examples of small synaptic vesicles (black arrows) in the bouton, a synapse between the bouton and soma (arrowheads), and an endocytic profile within the soma (white arrow). I) Two non-consecutive 50 nm sections of the central amygdala labeled with a probe for somatostatin mRNA (left) or antibodies to GABA and VGluT2 (right). A cell containing FISH signal and surrounded by punctate GABA signal is boxed, with the IF overlaid on a 2500X TEM image of the adjacent section. J) The area in the small box in the left panel of (I) imaged at 10,000X. The left panel shows an axonal bouton (white dashed line) overlaid with IF signal for GABA, and the right panel shows the EM image alone with a synapse between the bouton and the soma (arrowhead) and examples of small synaptic vesicles (arrows) indicated. The leftmost vesicle is docked at the active zone. Scale bar = 2 µm in (A-C), 10 µm in (F, I left), 3 µm in (G top), 500 nm in (G bottom), and 200 nm in (H,J).

### Serial multiplexing of light and electron microscopy

Fluorescence microscopy is very effective at revealing the spatial distribution of target molecules but offers little information about cellular context. Detailed subcellular structure can only be visualized by EM, and the option to integrate EM into an ultraplex workflow is therefore essential for certain applications. After modifying our embedding protocol to improve preservation of ultrastructural morphology, we first tested the use of cross-modal serial multiplexing to resolve ambiguous patterns of fluorescence labeling. In the central amygdala, double-label IF for calcitonin gene-related peptide (CGRP) and VGluT2 produces a striking pattern of dense, partially overlapping signal that encircles some nuclei (Figure 5F), suggesting that CGRP and VGluT2 colocalize in axons that form synapses onto neuronal cell bodies. EM imaging of adjacent sections confirmed that this is indeed the case. IF signal localized to large axonal boutons in direct apposition to cell bodies (Figure 5G), and these boutons formed synapses and contained both large dense-core vesicles (LDCVs) and small synaptic vesicles (Figure 5H). The two types of vesicles were distributed in semi-segregated pools, consistent with the partial overlap between VGluT2 and CGRP signal. When IF and FISH were applied across serial sections, we noted that cells containing somatostatin mRNA were not contacted by CGRP (not shown) or VGluT2, but did appear to have adjacent profiles filled with GABA signal (Figure 5I). Performing EM and IF for GABA on adjacent sections confirmed that the GABA signal corresponded to presynaptic vesicle pools in axons forming synapses onto somatostatin cell bodies (Figure 5J).

### Cross-modal ultraplexing for single-bouton profiling

Axonal boutons span multiple 50 nm sections, so it should be possible to profile single boutons across sections similarly to single cells (Figure 2E). As a test of single-bouton profiling, EM and IF were interleaved on a set of eight serial 50 nm sections of central amygdala. Each IF section was labeled with two of four antibodies – VGluT1, VGluT2, somatostatin, and VIP – and images were taken at the boundary between the medial and lateral subdivisions (Figure 6A) where signal for all four antibodies is intermingled. EM and IF images were aligned using large landmarks such as nuclei and blood vessels (Figure 6B), after which IF puncta could be assigned to boutons in the EM images (Figure 6C). Boutons could be easily followed across single-section skips so that morphology and molecular signal could be compared (Figure 6D). IF signal was generally found within the EM boundaries of a bouton on at least one adjacent section. As expected, VGluT1 and VGluT2 signal corresponded to pools of small synaptic vesicles and somatostatin and VIP to large dense-core vesicles (Figure 6E). Consistent with our observation of a significant fraction of somatostatin IF signal overlapping with GABA signal in the neuropil (Figure 4E-F), boutons labeled for somatostatin contained pools of small synaptic vesicles intermingled with large dense-core vesicles. Boutons labeled for VIP tended to contain only dense-core vesicles, so it is possible that VIP axons also contain other vesicle types in separate boutons. More than eight sections may be necessary to answer this question.

**Figure 6.**
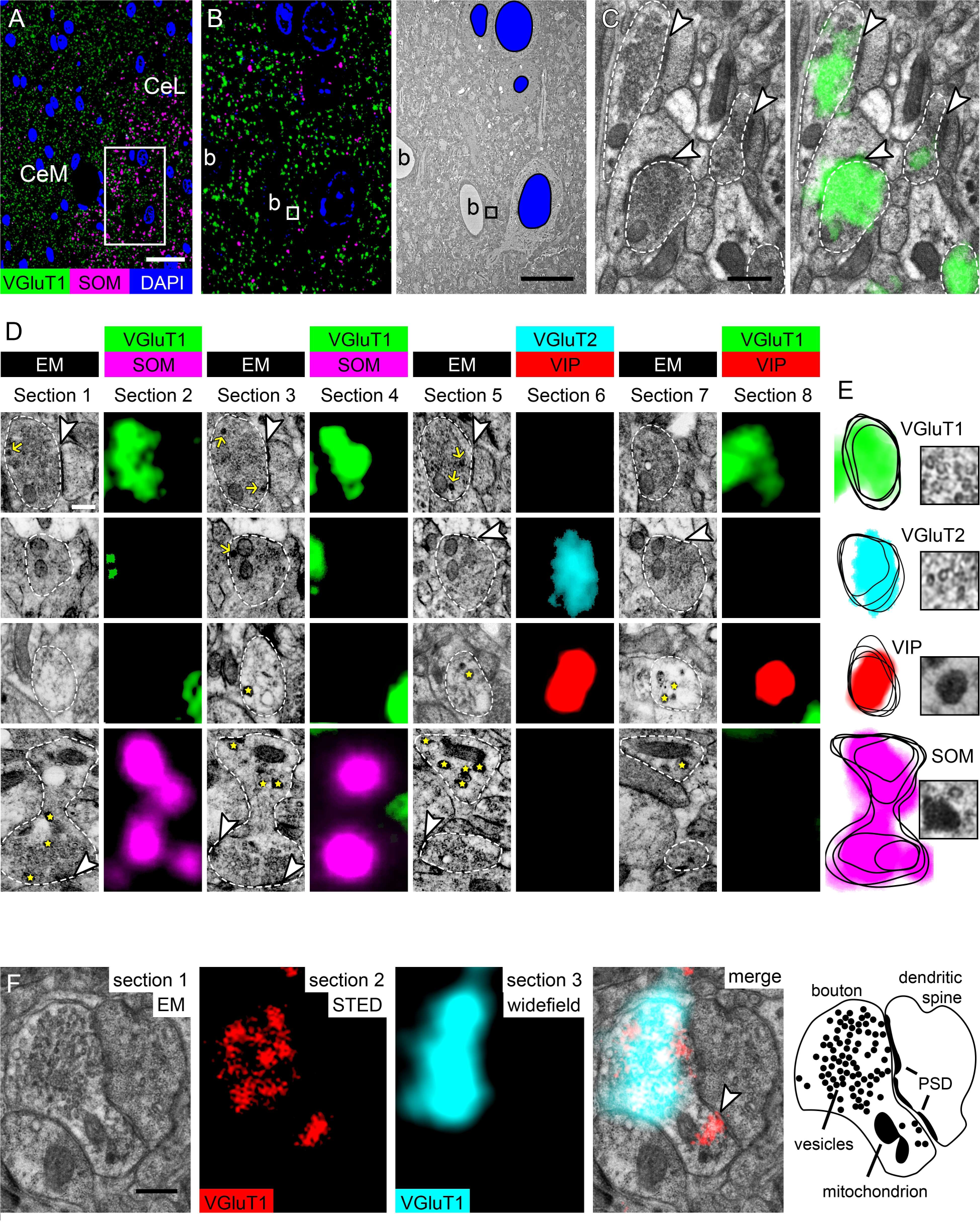
Multi-modal ultraplex microscopy at the single-bouton level. A) Immunofluorescence for VGluT1 and somatostatin at the boundary between the medial and lateral divisions of the central amygdala imaged at 20X. B) The boxed region in (A) imaged at 100X (left) and the same region on an adjacent section imaged by TEM at 600X (right) with nuclei (blue fill) and blood vessels (b). C) The boxed region in (B) imaged at 10,000X by TEM (left) and overlaid with the 100X IF image (right). Dashed outlines indicate axonal boutons, arrowheads indicate synapses onto dendritic spines. D) Sequence of immunofluorescence and EM across eight serial 50 nm sections. Each row shows a different bouton (white outlines in EM images). Synapses (arrowheads), small dense core vesicles (yellow arrows), and large dense-core vesicles (yellow stars) are visible in some EM images. E) Overlay of EM outlines and IF signal within each of the four boutons in (D), alongside a 3X enlargement of vesicles from an EM image. F) A synapse between a dendritic spine and an axonal bouton on three consecutive 50 nm sections. Section 1 was imaged at 10,000X by TEM and sections 2 and 3 were labeled with the same primary antibody to VGluT1 and imaged by STED and widefield fluorescence, respectively. The arrowhead in the merged image indicates a region of the synapse where the synaptic cleft, docked vesicles, and the postsynaptic density (PSD) are easily visible. The far right panel shows an outline of the structures in the EM image. Scale bar = 25 µm in (A), 10 µm in (B), 300 nm in (C), 200 nm in (D), and 250 nm in (F).

Super-resolution fluorescence microscopy lacks the ease and efficiency of widefield imaging, but its improved resolution may be useful in some circumstances. As a test of tri-modal ultraplexing, we performed TEM, stimulated emission depletion (STED), and widefield epifluorescence microscopy on three consecutive sections, with the same antibody to VGluT1 applied to the STED and widefield sections. Signal for VGluT1 mapped onto EM-verified axonal boutons, with STED signal hewing more closely to the contours of vesicle pools than widefield (Figure 6F).

### Correlative confocal and ultraplex microscopy

We imaged fluorescent proteins on 50 nm sections that had been rehydrated and coverslipped in an aqueous mounting medium (Figure 4), but if fluorophores could be imaged in the intact block it would be possible to perform correlative confocal and ultraplex microscopy. This could be useful for locating cells after live imaging, for example, or visualizing cell morphology without extensive reconstruction of serial ultrathin sections. To test confocal imaging in plastic-embedded tissue, injection site blocks were trimmed to expose tissue at the surface and mounted on glass coverslips with an aqueous mounting medium containing DAPI. In the thalamus, tdTomato-expressing cells, GFP-filled axons from the cortex, and DAPI-stained nuclei were visible by confocal (Figure 7A). Both fluorescent proteins could be imaged to a depth of at least 30 μm, but DAPI only penetrated about 5 μm into the blocks (Figure 7B).

**Figure 7.**
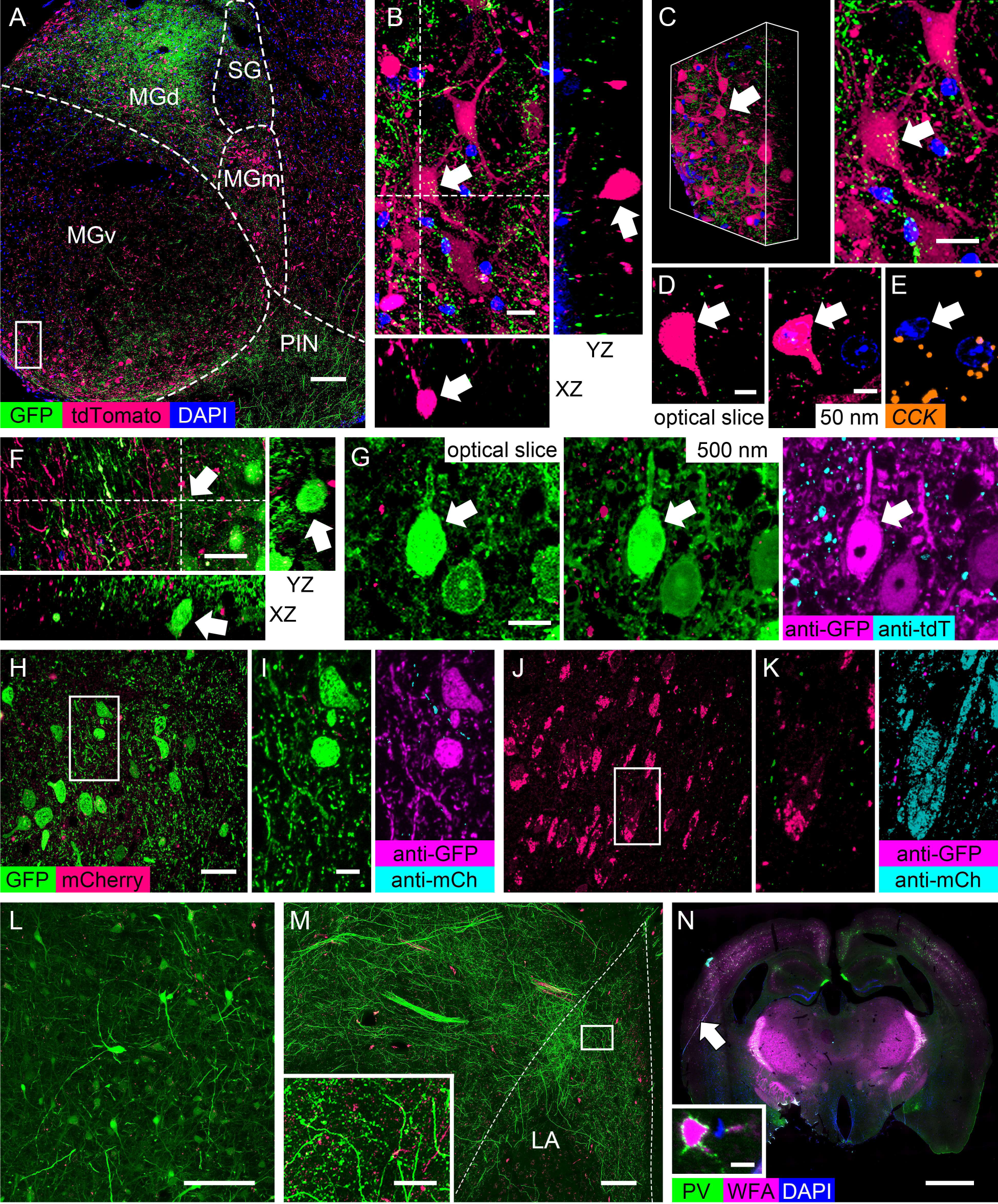
Correlative confocal and ultraplex microscopy. A) Confocal image of plastic-embedded auditory thalamus expressing tdTomato with GFP-expressing axons from the auditory cortex. Maximum intensity projection of 93 0.5 µm optical slides imaged at 63X is shown. B-E) Multiple views of the same tdTomato-expressing cell (arrows). B) Enlarged view of the boxed area in (A) with XZ and YZ planes orthogonal to the dashed lines. C) Projection of the image stack (left) and enlarged maximum intensity projection (right). D) A single optical section (left) and a 60X image of a 50 nm section collected after confocal imaging and imaged by widefield (right). E) A 50 nm section adjacent to the one in (D) labeled with a probe for CCK mRNA. F-H) Multiple views of a GFP-expressing cell (arrows) in the auditory cortex. F) Maximum intensity projection and orthogonal planes through 63 0.5 µm optical slices the intact plastic block imaged at 63X. G) A single optical section (left), native fluorescence on a 0.5 µm section (center), and antibody labeling for GFP and tdTomato on the same 0.5 µm section. H-K) Native fluorescence on 50 nm sections from plastic blocks stored for 18 months under ambient conditions, imaged at 60X. H) GFP-expressing cells in the auditory thalamus with mCherry-expressing axons from the auditory cortex. I) Enlarged view of box in (H) showing native fluorescence of GFP and mCherry (left) and antibody labeling for the two fluorophores on an adjacent section (right). J) Cells in the auditory cortex expressing mCherry with GFP-expressing axons from the auditory thalamus. K) Enlarged view of the boxed region in (J), showing native fluorescence (left) and antibody labeling (right). L) Confocal image of plastic-embedded auditory cortex after 18 months of storage, showing GFP-expressing cells and mCherry-expressing axons. M) Confocal image of plastic-embedded amygdala from the same brain as the block in (L), showing GFP-expressing axons from the thalamus and mCherry axons from the cortex. Inset: Enlarged view of the boxed region in (M). N) IF for parvalbumin (PV) and staining with wisteria floribunda agglutinin (WFA) on a 100 nm section of mouse brain imaged at 20X. Inset: Enlarged view of the cell indicated by the arrow. MGm, MGv, MGd: medial, ventral, and dorsal divisions of the medial geniculate nucleus, respectively; PIN: posterior intralaminar nucleus; SG: suprageniculate nucleus; Ce: central amygdala; LA: lateral amygdala. Scale bar = 100 μm in (A,M), 10 μm in (B, C, G, and N inset), 5 μm in (D,E,I,K), 20 μm in (F, M inset), 15 μm in (H, J), 1 mm in (N).

After peeling off the mounting medium, blocks were ultrathin sectioned for labeling. Individual neurons in the confocal volume (Figure 7C) could be identified by matching ultrathin sections to single optical slices (Figure 7D). Native fluorescence in the ultrathin sections was quenched by the FISH protocol (Figure 7E), so serial sections without FISH were necessary to identify FISH-labeled cells in the confocal volume. GFP-expressing neurons and thalamic axons containing tdTomato in the auditory cortex were also visible by confocal (Figure 7F). To compare optical and physical sections, 0.5 µm optical sections were acquired at 63X by confocal and 0.5 µm sections were then cut and imaged by widefield at 60X. The same cells and processes could be seen in both imaging modalities, and antibodies to GFP and tdTomato on the 0.5 µm section indicated that the native fluorophore signals were preserved (Figure 7G).

To assess the longevity of fluorescent proteins in plastic-embedded samples, we examined blocks from rats expressing GFP in the auditory thalamus and mCherry in the auditory cortex. Neurons filled with GFP and axons containing mCherry were visible in 50 nm sections of auditory thalamus (Figure 7H), and antibody labeling for both proteins indicated that the GFP signal was largely intact (Figure 7I). In the auditory cortex, mCherry was visible in neurons but signal was uneven (Figure 7J). Antibody labeling demonstrated that while GFP signal was complete in axons, a substantial amount of mCherry signal had faded (Figure 7K). Confocal imaging of an intact block of thalamus found that GFP-filled cells were readily visible while mCherry axons were faint (Figure 7L), and in an amygdala block dense GFP axons from the thalamus were easily visible and mCherry axons from the cortex were detectable but had weaker signal (Figure 7M).

### Large-format sections

Coronal sections of rodent brains are a standard format for IF and FISH but ultrathin sections for EM are much smaller, typically no more than 2 – 3 mm on a side. We embedded full coronal sections of adult mouse brain and found that they could be cut reliably at 100 nm for labeling (Figure 7N).

## Discussion

No single microscopy method can visualize tissue in complete molecular and structural detail across the full range of scales and resolutions, and the potential for combining multiple techniques is constrained by incompatibilities between sample preparation methods, staining protocols, and labeling reagents. The limited breadth of available imaging options is a particularly important problem in neurobiology due to the dramatic scope of complexity in brain structure. Cells with heterogeneous morphology and transcriptomes intermingle in brain tissue^1,2^, and temporal and spatial regulation of protein expression throughout large, highly polarized cells mean that RNA transcripts and their protein products can appear at different times and even in different brain regions^3,12,13^. A means of simultaneously labeling endogenous RNA and proteins with sub-cellular resolution is thus essential for understanding the molecular organization of the brain. Ultraplex microscopy enables otherwise impossible combinations of microscopy techniques to be applied to a single sample using the complementary advantages of serial multiplexing, ultrathin sectioning, and reversible embedding.

By dividing samples into separate sections for parallel, independent experiments, serial multiplexing eliminates constraints on multiplexed imaging imposed by the number of imaging channels and by interference between labeling protocols. The latter can include cross-reactivity between antibodies, digestion of antibody targets by proteases used in FISH, degradation of RNA by antigen retrieval treatments, and distortion of tissue structure by detergents used to permeabilize tissue for labeling^11,15^. The mirror technique is a form of serial multiplexing that consists of applying different antibodies to adjacent fixed tissue sections and examining the facing surfaces for cells bisected by the sectioning plane^16–18^. Sections must be thicker than the diameter of a neuron to avoid deformation^16^, which means that many cells are not exposed at the surface, a cut cell can appear on no more than two sections, and only one face per section can be analyzed. The effort of tracking facing section surfaces is not worthwhile unless few section pairs are needed, as in serial multiplexing of EM with light-level immunolabeling^18^.

In contrast to fixed tissue, samples embedded in EM resin can be cut into uniform serial sections as thin as 30 nm while retaining intact subcellular morphology^19–21^, which increases serial multiplexing potential by two orders of magnitude over the mirror technique. The two types of antigens most amenable to immunolabeling on conventional EM sections – peptide hormones and amino acid neurotransmitters^26^ – have already been used for serial multiplexing at the EM level^23,24^, including in large-scale volume reconstructions^22,25^. EM resins are crosslinked to entrap tissue in a dense three-dimensional matrix, and the difficulty of antibody labeling on standard EM sections is attributed to the inability of reagents, especially those > 300 D, to enter the dense tissue-resin matrix, as well as by exclusion of hydrophilic reagents by nonpolar resin components and, in the case of epoxy resins, denaturation of tissue molecules by reaction with the resin itself^28^. Antigenicity can be improved on epoxy resin sections by breaking the covalent bonds with aggressive etching agents^27–29,35^ or by replacing epoxy resins with acrylic resins such as LR White, which is argued to be more compatible with immunoreagents than other resins because it is hydrophilic^56^. High concentrations of antibodies are needed for labeling on any EM resin^32–34^, however, and specialized screening is needed for antibodies compatible with LR White^36^. Etched epoxy resin performs better in IF than acrylic resins^32^, but the few reports of successful RNA labeling on resin – mainly of high-abundance targets such as poly-A, viral, or ribosomal RNA – used hydrophilic acrylic resins^57–64^.

Reversible embedding allows sections to be labeled without interference from the embedding medium or damage from its removal. We formulated a non-crosslinked, acetone-soluble embedding plastic for cutting serial ultrathin sections that could be used for routine labeling procedures without the trade-offs imposed by EM resins. Deplasticized ultrathin sections proved to have advantages over both resin sections and conventional fixed and fresh-frozen sections, in fact. IF labeling could be performed using the same antibodies that we use on fixed tissue sections at the same or even lower concentrations, but with an incubation time of one hour instead of overnight. IF signal on deplasticized sections could often be further enhanced by heat-induced antigen retrieval, but this had to be omitted in comparisons with plastic and LR White sections because it caused them to detach from glass substrates.

Ultrathin sections have another crucial, unique advantage in multiplexing that became especially clear when deplasticized sections were used: because ultrathin sections are essentially two-dimensional, reagents do not need to penetrate into the tissue to generate detectable signal. This is critical because reagent penetration in fixed sections is a function of primary fixation, and primary fixation is a major source of protocol incompatibility that cannot be addressed by serial multiplexing because it occurs before sectioning. Acceptable fixation varies widely among labeling and imaging methods. Standard EM protocols start with strong glutaraldehyde fixation to maximize retention of molecules and tissue structure during subsequent processing, and glutaraldehyde fixation is also essential for immunolabeling of some antigens, including GABA and other small molecules ^23,26^. Milder fixation with formaldehyde is used for most other antigens, and FISH protocols often use fresh-frozen tissue to minimize chemical fixation. Glutaraldehyde is generally considered incompatible with most molecular labeling because, unlike formaldehyde, it forms dense, irreversible crosslinks that inhibit labeling reagents from reaching their targets^65^. Combining protocols with differing fixation needs, particularly where glutaraldehyde is concerned, requires significant compromise^15^. In pilot tests we found that glutaraldehyde fixation improved EM morphology and was necessary for immunolabeling amino acids including GABA, as expected, but that it did not qualitatively affect FISH or most IF labeling on deplasticized ultrathin sections. This enabled the unusual combination of FISH with EM, and IF for GABA with other antigens and FISH. In contrast, we were unable to use glutaraldehyde with conventional preparations because it abolished FISH labeling and reduced IF signal. The addition of uranyl acetate fixation greatly improved EM preservation and did not have any apparent effect on IF or FISH, which was not surprising as uranyl acetate is reported to improve immunolabeling on EM sections^66^.

Ultraplex microscopy does not require any customized reagents, protocols, materials, or equipment, and although the workflow itself is novel, the procedures are not. We used routine labeling reagents at routine concentrations on our sections, and imaged them on standard microscopes without modifications. Our embedding formula is composed of readily available, inexpensive components and the embedding steps will be familiar to anyone who has worked with UV-cured embedding media. We describe two embedding protocols which may be of use in different contexts. For optimal preservation of EM structure, we used high-pressure freezing, freeze-substitution, and cryoembedding methods^67,68^ similar to those used for Lowicryl^69,70^. These samples have some autofluorescence due to glutaraldehyde and uranyl acetate, and although this did not interfere with IF or FISH signal, imaging of fluorescent proteins without antibody amplification benefited from a reduced concentration of uranyl acetate. Instruments for cryoembedding may not be available to non-electron microscopists, and our general protocol calls for UV curing at 0°, a temperature that avoids heat damage to fluorescent proteins and tissue molecules yet can be maintained without expensive equipment.

The need for serial ultrathin sectioning would seem to be a barrier to routine use of ultraplex microscopy, given that expertise in ultrathin sectioning is generally confined to electron microscopists and serial sectioning is even further specialized, but we do not expect this to be the case. Serial sectioning for volume EM requires advanced skills^19,20^ and equipment^21^ because hundreds of sections must be collected on EM substrates with little to no tolerance for section loss. Serial ultrathin sections are also used in array tomography, a technique that uses LR White for volume reconstruction of high-resolution IF images^33,34^. Sectioning for array tomography is even more challenging than serial EM sectioning because the fragile hydrophilic sections are held together with glue and a special knife must be purchased^71^. Our embedding formula is easy to serial section and the sections are even robust enough to collect with a brush. In addition, the loss of a few sections or shuffled section order would not affect the results in many applications of ultraplex microscopy – single-cell profiling for example, or within section analysis.

The sample format and workflow we have described allows individual samples to be imaged in greater breadth and detail than is possible with any other microscopy method. It is unique in allowing samples to be fixed strongly enough to retain EM morphology and small neurotransmitters without impairing detection of other antigens or RNA targets, and it is the only microscopy approach that essentially aliquots imaging fields for separate experiments. Its versatility makes it valuable in many applications beyond the ones we tested. Because RNA is preserved, it may be possible to perform *in situ* sequencing on 50 nm sections^6^, for example, and the clarity of the embedding resin could allow for light-sheet imaging^72,73^. In this study we have focused on brain tissue, but the technique should be useful in a wide range of experimental systems where multiplexed imaging or sensitive high-resolution labeling are needed.

## Materials and Methods

### Subjects

Adult male (n = 8) and female (n = 1) Sprague-Dawley rats (Hilltop Lab Animals, Scottdale, PA or Charles River Laboratories, Wilmington, MA) aged 8-12 weeks and one adult male C57BL/6 mouse (Jackson Laboratories, Farmington, CT) were pair-housed on a reverse 12-hour light cycle in ventilated cages with *ad libitum* food and water. All animal procedures were approved by the University of Connecticut Institutional Animal Care and Use Committee.

### Virus Injections

Animals were anesthetized with ketamine (100 mg/kg) and xylazine (10 mg/kg) and placed into a stereotaxic frame (Kopf Instruments, Tujunga, CA). AAV vectors were infused through cannulae using a syringe pump (KD Scientific) at a rate of 2.65 mL/min, for a total volume of 0.3 μL in the auditory thalamus (AP:3.2, ML:3.0, DV:3.3 mm from the interaural center) and 0.5-0.6 μL in the auditory cortex (AP:3.8, ML:6.8, DV:3.2 mm from the interaural center). Viral vectors were purchased from Addgene and used as received. Titers were AAV8-hSyn-EGFP (≥ 5×10¹² vg/mL), AAV8-hSyn-mCherry (≥ 7×10¹² vg/mL), and a 1:1 mixture of AAV1-hSyn-Cre-WPRE-hGH (≥ 7×10¹² vg/mL) and AAV8-CAG-FLEX-tdTomato (≥ 1×10¹³ vg/mL).

### Tissue Collection

Animals were deeply anesthetized with chloral hydrate (750 mg/kg). For collection of fixed tissue, animals were perfused transcardially with 500 mL of 4% freshly depolymerized paraformaldehyde and 1% glutaraldehyde in 0.1M phosphate buffer, pH 7.4. Glutaraldehyde was omitted from the fixative for brains expressing fluorescent proteins to minimize autofluorescence. Brains were immersed in the perfusion fixative for one hour at room temperature, and brains perfused with only paraformaldehyde remained immersed in the perfusion fixative overnight at 4°C. Brains were rinsed in the perfusion buffer and sectioned coronally on a vibrating microtome (Leica Biosystems, Wetzlar, Germany). The region of interest was then isolated with a 2 mm biopsy punch. For collection of fresh-frozen tissue, anesthetized animals were decapitated and the brain was quickly frozen on dry ice, surrounded with optimal cutting temperature (OCT) medium, and stored at - 80°C.

### Embedding formula

To prepare the embedding formula, methyl methacrylate and *n*-butyl methacrylate combined in a 2:3 ratio and bubbled with dry nitrogen in a glass container for 15 minutes to mix. Benzoin methyl ether is then added for a final concentration of 0.5% and the mixture is bubbled for another 15 minutes. The embedding mixture can be used immediately, or stored up to a week at -20°C and bubbled with nitrogen again before use. All embedding reagents were obtained from Electron Microscopy Sciences (Hatfield, PA).

### Embedding for light microscopy

All dehydration and infiltration steps were performed in a laboratory microwave (Biowave Pro; Ted Pella, Redding, CA) with the power set to 750W. Biopsy punch samples 200 – 400 μm thick were dehydrated in an ascending series of ethanol dilutions in water (50%, 70%, 90%, and 2 x 100%) containing 1 mM dithiothreitol (DTT; ThermoFisher Scientific) for 40 seconds per step with the temperature restriction at 37°C. For infiltration, samples were transferred to a 1:1 dilution of the embedding mixture and ethanol containing 1 mM DTT, followed by two changes of pure embedding mixture with 10 mM DTT. Each infiltration step was carried out for 15 minutes with the temperature restriction at 45°. Samples were transferred into gelatin capsules filled with fresh embedding mixture with 10 mM DTT and cured for 48 hours under a UV-A lamp (380 nm LED) at 0°C in the chamber of an automated freeze-substitution device (AFS2; Leica Microsystems, Wetzlar, Germany).

### Embedding in LR White

LR White resin (Electron Microscopy Sciences) was used according to the manufacturer’s instructions. The resin was purchased uncatalyzed and the catalyst was added and mixed for 24 hours before use. Samples were dehydrated and infiltrated as described above, then transferred into gelatin capsules of fresh resin and cured for 48 hours at 60°C. No DTT was added to the samples.

### Embedding for electron microscopy

Biopsy punches 100 μm in thickness were placed in aluminum carriers (well dimensions: 2 mm diameter x 100 μm depth) with 20% polyvinylpyrrolidone (EMD Millipore Corp., Burlington, MA) in water as a filler, then frozen in a Wohlwend Compact 3 high-pressure freezer (Technotrade International, Manchester, NH) and stored under liquid nitrogen until further processing. To prepare for embedding, samples were freeze-substituted in acetone containing 1% uranyl acetate (Structure Probe, Inc., West Chester, PA) for 48 hours at -90°C in the automated freeze-substitution device, then rinsed with pure acetone before warming to -35°C at 5°C per hour. Infiltration was performed at -35°C by the following schedule: 1:1 and 2:1 embedding mixture:acetone with 1 mM DTT for 3 hours each; pure embedding mixture with 10 mM DTT overnight, then for one hour. Samples were then transferred into polyethylene flat embedding capsules (Electron Microscopy Sciences) filled with fresh resin and cured under a UV-A lamp at -35°C for 48 hours and warmed to 20°C at 5°C per hour with the UV light on.

### Sectioning

Blocks were hand-trimmed with razor blades and sectioned on a Leica UC7 ultramicrotome with a clearance angle of 4° and cutting speed of 1 mm/s. For light microscopy imaging, a small paintbrush or a perfect loop was used to transfer sections on #1.5 glass coverslips for fluorescence imaging or glass slides for histology. Sections were allowed to air dry before further use. For experiments involving LR White or staining of sections with intact plastic, coverslips or slides were coated with gelatin to prevent sections falling off during labeling. For electron microscopy imaging, sections were collected on slot grids coated with pioloform (Ted Pella, Inc.). All other sectioning supplies were obtained from Electron Microscopy Sciences (Hatfield, PA).

### Plastic removal

Sections were deplasticized by incubating in acetone for 10 minutes, then rehydrated in a descending series of ethanol dilutions in water (100%, 70% 50%) for 1 minute each, followed by a 2 minute water rinse.

### Histological staining

For toluidine blue staining, a solution of 0.5% toluidine blue and 0.5% sodium tetraborate was applied to sections mounted on coverslips. Coverslips were heated for about 2 minutes on a 60°C hotplate, rinsed thoroughly with distilled water, allowed to dry, and coverslipped with DPX (Electron Microscopy Sciences). For Hematoxylin and Eosin (H&E) staining, sections were dehydrated with acetone for 10 minutes, then rehydrated through a series of descending ethanol dilutions in water (100%, 70% 50%) for 1 minute each with a final rinse in water for 2 minutes. Reagents in the staining kit were used as received, except that hematoxylin was filtered through Whatman #1 filter paper before use. Hematoxylin was applied to sections for 8 minutes, and after water rinses, five quick dips in 0.25% acid alcohol (2 ml of 37% HCl in 300 ml of 50% ethanol in water) and 20 dips on bluing reagent. Sections were thoroughly rinsed in distilled water for 10 minutes, then quickly incubated in 95% ethanol and Eosin Y for 1 minute prior to dehydration with ascending ethanol series in water (95%, 2x-100%) for 1 minute each. Sections were allowed to dry, then coverslipped with DPX.

### Immunofluorescence on ultrathin sections

Sections were blocked for 30 minutes in PBS (9% sodium chloride in 0.01M sodium phosphate, pH 7.4) containing 1% bovine serum albumin (Jackson ImmunoResearch Laboratories Inc. West Grove, PA) and either 0.1% Tween-20 or 0.3% Triton X-100. Sections were incubated for 1 hour in primary antibodies diluted in blocking buffer, then rinsed twice for 2 minutes in PBS containing 0.1% Tween 20 (PBST) and incubated in secondary antibodies diluted in blocking buffer for 30 minutes. All antibodies and concentrations are listed in Supplemental Table 1. Sections were rinsed twice in PBST, then rinsed once in PBS containing 2 μg/ml DAPI, and finally rinsed in PBS and mounted in ProlongGold (Invitrogen). All incubations were performed at room temperature in a humidified plastic tray to prevent sections drying.

### Heat-induced antigen retrieval

Antigen retrieval was performed in a pressure cooker (Instant Pot, Inc.) in 0.1M sodium citrate, pH 6.3. Sections were placed in preheated buffer, held under pressure for 10 minutes, then immediately removed and rinsed twice in 0.01M phosphate saline buffer, pH 7.4 with 0.1% Tween-20 (PBST) at room temperature.

### Cyclic labeling

Sections were imaged within an hour of mounting and the coverslip was removed by sliding it sideways while the mounting medium was still wet. Antibodies were eluted using a modified version of a protocol used by Holderith and colleagues^32^. Sections were incubated in 1% sodium dodecyl sulfate in PBS for 10 minutes in a pressure cooker, as for antigen retrieval, then rinsed in PBS before a second round of labeling.

### Immunofluorescence on fixed sections

Free-floating 40 µm vibratome sections of 4% paraformaldehyde fixed brains were blocked for 1 hour in 0.01M PBS containing 1% BSA, then incubated overnight in primary antibodies diluted in blocking buffer at room temperature with agitation. After three 5 minute rinses in PBS, the sections were incubated with secondary antibodies diluted in blocking buffer for 1 hour. Sections were rinsed in PBS with 2 µg/ml DAPI, mounted on glass slides, and coverslipped with ProlongGold.

### Fluorescence in situ hybridization

For detection of poly-A RNA, sections were rinsed twice with sterile PBST and incubated for 5 minutes in a solution of 1µg/ml of proteinase K diluted in PBS. After PBST rinses, sections were incubated in acetic anhydride solution (100mM TEA pH 8.0, 0.24% acetic anhydride) for 5 minutes then rinsed in PBST and incubated for 1 hour at 60°C in prewarmed hybridization buffer (10mM Tris pH 7.4, 600mM NaCl, 1mM EDTA, 0.25% SDS, 10% dextran sulfate, 1X Denhardt’s solution, 200µg/ml yeast tRNA, 50% deionized formamide in sterile water). Probe (2µM) (20-bp poly-A-3’biotin at 2 μm; Integrated DNA Technologies, Coralville, IA) was diluted in hybridization buffer and incubated for 2 hours at 60°C. Sections were rinsed for two minutes in saline sodium citrate buffer (SSC) at 60°C (1X + 50% deionized formamide (EMD Millipore Corp., Burlington, MA), 2X, and 2x 0.2X). Sections were then rinsed once in PBST followed by two rinses in 1X maleic acid buffer containing 0.1% Tween-20 (MABT). Sections were blocked (3% Heat-inactivated Sheep Serum (HISS), 0.1% blocking reagent (EMD Millipore Corp., Burlington, MA) in MABT for 10 minutes at room temperature, then incubated in streptavidin-AlexaFluor 647 diluted in blocking buffer for 15 minutes at 4°C. Rinses in MABT followed a 2-minute incubation in 4% paraformaldehyde in PBS. Sections were rinsed in PBS with 2 µg/ml DAPI, coverslipped with ProlongGold Antifade Mountant, and dried overnight at room temperature.

The RNAscope Multiplex Fluorescent Reagent Kit v2 (Advance Cell Diagnostics Inc., Newark, CA.) was used according to the manufacturer’s instructions. Sections were dehydrated and rehydrated as previously mentioned (see Immunofluorescence). After rehydration, the area containing sections was encircled with PAP Pen (Vector Laboratories, Newark, CA. Catalog no. H-4000) to keep all solutions on the sections. Sections were incubated for 10 minutes with hydrogen peroxide. They were then incubated for 30 minutes at room temperature using protease III. Other proteases and antigen retrieval solutions were tested with no optimal results. Probes were incubated for 2 hours, followed by amplifiers (AMP 1, 2, and 3) and HRP-C1 for all probes. The signal was detected using TSA Vivid 650 diluted at 1:500 in the provided 1X amplification buffer. Sections were incubated for 30 seconds with DAPI (provided in the kit), coverslipped with ProlongGold Antifade Mountant and dried overnight at room temperature. All incubations used the HybEZ^TM^ II Oven at 40°C. Rinses were done using EasyDip Slide Staining Jars (Fisher Scientific. Catalog No. 22-038-491).

For fresh-frozen sections, brains stored at -80°C were sectioned at 5 μm on a cryostat and sections were mounted on SuperFrost Plus Gold microscope slides (Fisher Scientific, Pittsburg, PA). Prior to RNAscope, sections were fixed with 4% paraformaldehyde at 4°C for 15 minutes and dehydrated in an ascending series of ethanol dilutions in water (50%, 70%, 100%) for 5 minutes at room temperature. RNAscope was performed as for ultrathin sections.

### Brightfield microscopy

Images were taken on a Leica Thunder Imager with a DMC5400 CMOS camera (Leica, Deerfield, IL) using Leica 10x/.45 Plan Apo and 63x/1.40 Plan Apo lenses.

### Widefield fluorescence microscopy

Sections were imaged on either a Nikon Eclipse Ni-E microscope equipped with an Orca-Fusion Digital CMOS camera (Hamamatsu, Bridgewater, NJ), a Nikon Ti2-E microscope equipped with a Prime 95B sCMOS camera (Teledyne Photometrics, Tucson, AZ), or a Leica Thunder Imager with a DMC5400 CMOS camera (Leica, Deerfield, IL). Details of imaging instrumentation and settings are given in Supplemental Table 1.

### Confocal microscopy

Images were collected on a Leica SP8 confocal DMI6000 inverted microscope. For experiments using cryosections and vibratome sections a 40X/1.30 HC PL APO Oil CS2 objective lens was used. After microtome sectioning and exposing the tissue to the surface of the blocks, embedded 200 or 400 µm sections were mounted to a µ-Slide 1 Well Glass Bottom coverslip (Ibidi, Grafelfing, Germany) using ProlongGold Antifade Mountant with DAPI. Slides were allowed to dry overnight, and imaging was performed using a 20X/0.75 HC PL APO IMM CORR CS2 and a 63X/1.40 HC PL APO Oil CS2 objective lens.

### STED microscopy

Imaging was performed on a STED system (Abberior, Gottingen, Germany) with an Olympus IX83 microscope using a pulsed excitation laser of λ= 561 nm and a pulsed STED depletion laser of λ= 775 nm. Images were acquired with a 100X/1.4 UPlanSApo objective lens.

### Electron microscopy

Grids were imaged in a JEOL 1400 transmission EM with an AMT Nanosprint-43 Mark II digital camera (Advanced Microscopy Techniques, Woburn, MA) at an accelerating voltage of 120 kV.

### Colocalization and signal intensity analysis

Images were thresholded by subtracting the average pixel value of background areas between labeled structures. For comparisons between native fluorescence of reporter proteins and immunofluorescence signals, a gamma adjustment of 0.5 was first applied to further normalize background. Colocalization analysis was performed using the coloc2 plugin in FIJI^74^. Manders’ coefficients^75^, Pearson’s R, and integrated intensity were compared using ANOVA followed by a Bonferroni post hoc test when indicated.

### Image processing

Photoshop software (Adobe, Inc.) was used to align serial section images and to adjust display settings. Adjustments were applied uniformly across each image.

## Supporting information

Key Resources Table

## Acknowledgements

We are grateful to Chris O’Connell and Emery Ng for assistance with STED imaging and to Ethan Gasteyer, William P. Armstrong IV, and Monica S. Antony for assistance with some of the experiments and to Joshua Johansen and Mark Terasaki for thoughtful comments on the manuscript. This work was supported by NIH MH130472, NIH MH129269, NSF 2014862 to LO, NIH S10OD023618 to Christopher O’Connell, and NIH S10OD016435 to Akiko Nishiyama. This work used resources of the Advanced Light Microscopy Facility and Bioscience Electron Microscopy Laboratory at the University of Connecticut.

## Author contributions

JPG and LO designed the project; JPG, JO, ZD, GR, RT, and IC performed experiments; JPG and LO wrote the manuscript with input from all authors.

## Supplemental Figure Legends

**Figure S1.**
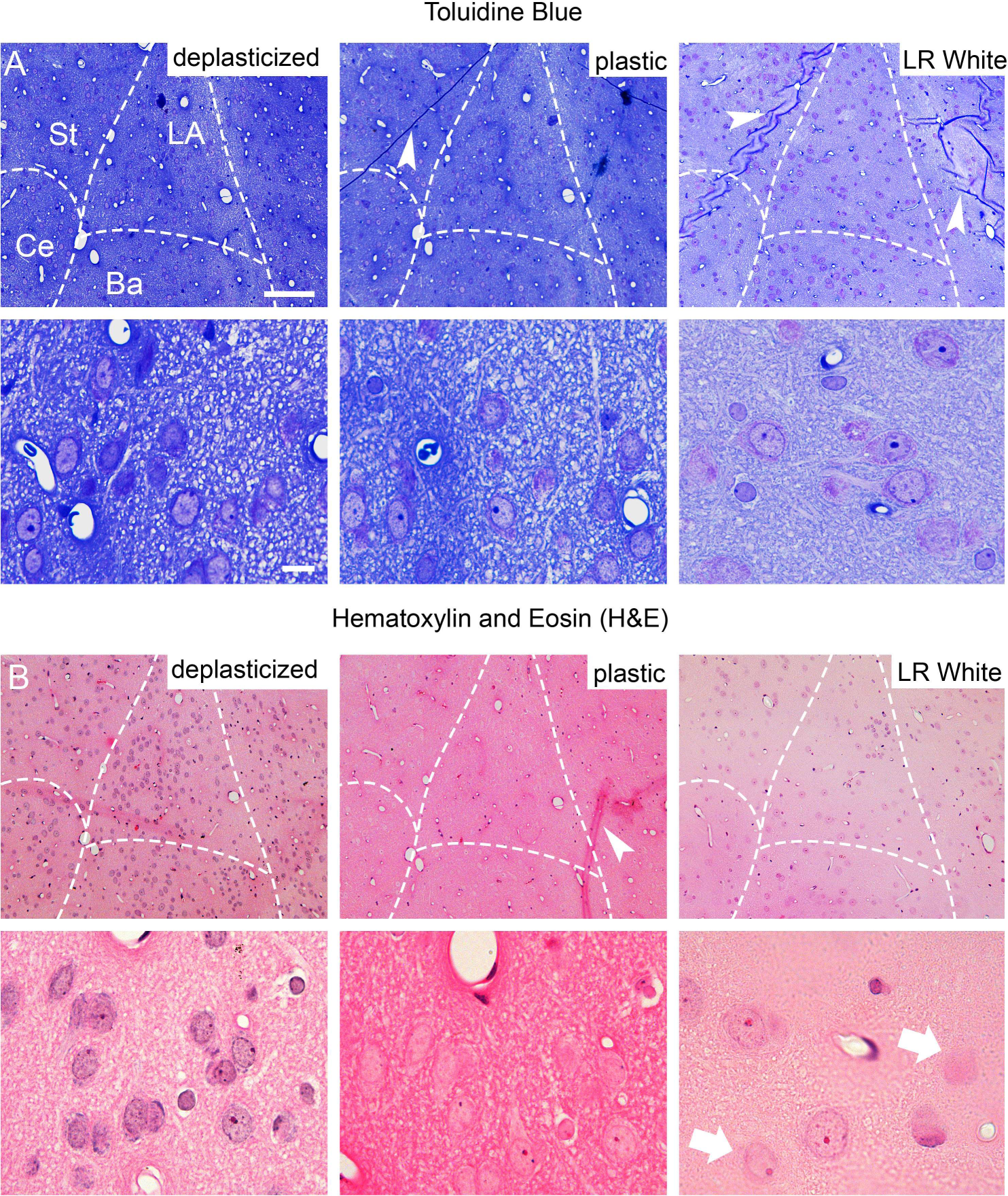
Brightfield imaging for histology. A) Toluidine blue staining on 1 µm sections of rat amygdala in deplasticized (left), plastic (center), and LR White (right) imaged at 10X. B) Hematoxylin and eosin staining (H&E) on 2 µm sections of rat amygdala in deplasticized (left), plastic (center) and LR White (right). Lower panels show areas from (A-B) imaged at 60X. Arrowheads show plastic sections folds. Arrows show blurry cells on uneven sections due to staining processing. Abbreviations as in Figure 1. Scale bar = 100 µm in (A-B, top panels), and 10 µm in lower panels.

**Figure S2.**
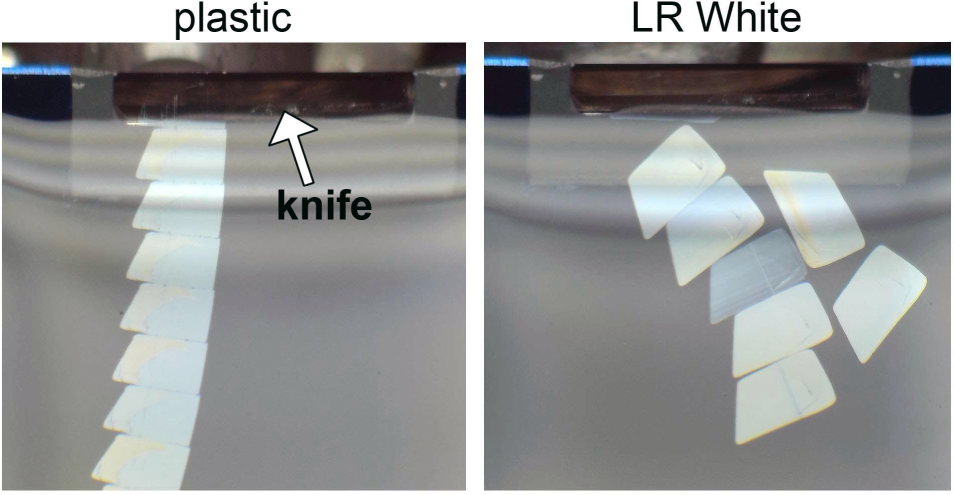
Serial ultrathin (50 nm) sections of brain tissue in plastic (left) and LR White resin floating in the boat of diamond knife. The knife is 8 mm wide.

**Figure S3.**
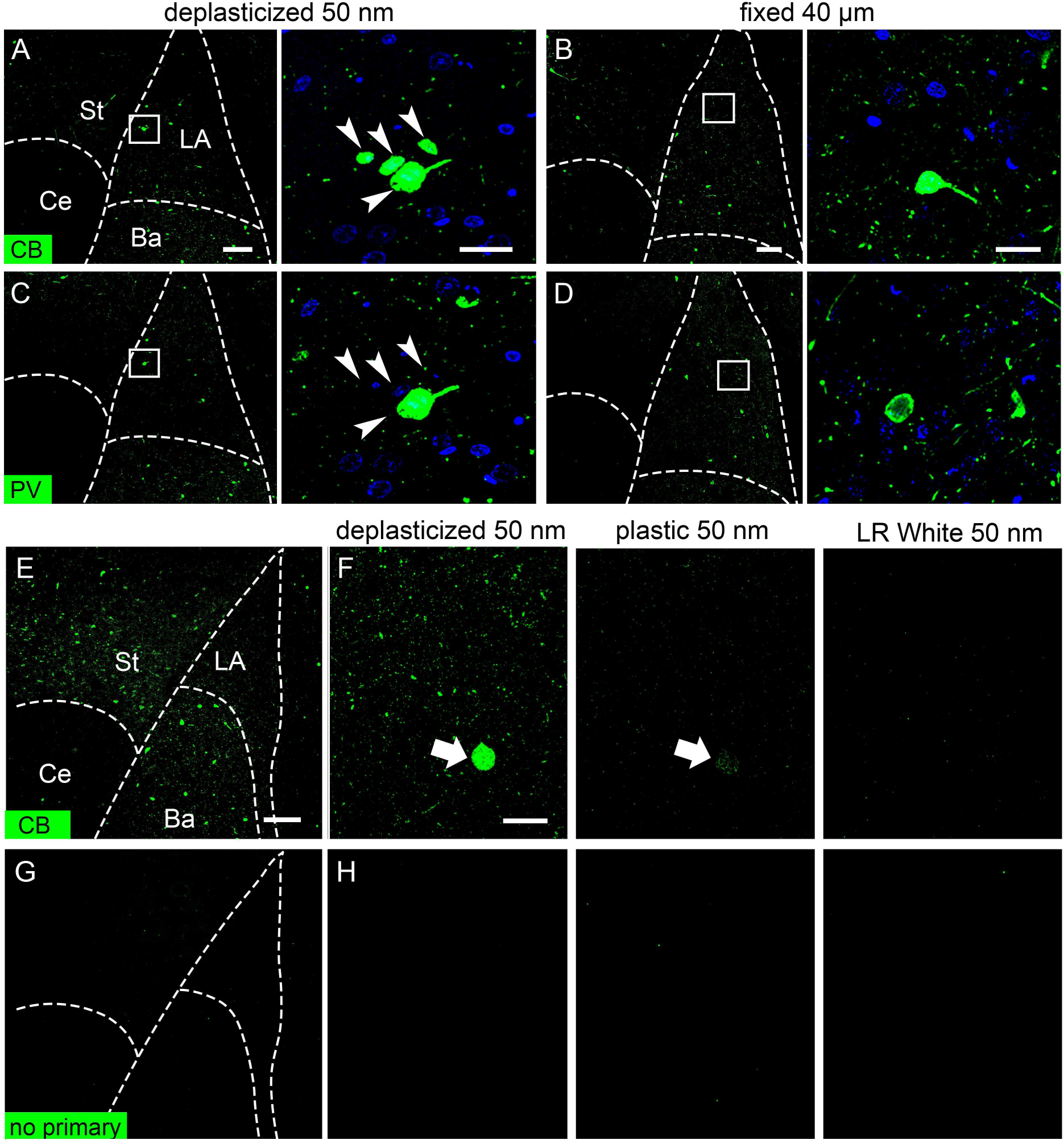
IF labeling on standard preparations and deplasticized ultrathin sections. A-D) Immunofluorescence for calbindin (A-B) and parvalbumin (C-D) on deplasticized 50 nm sections (A,C) or 40 µm vibratome sections of fixed rat amygdala (B,D). Ultrathin sections were imaged at 20X on a widefield microscope and single 0.5 µm optical sections of vibratome and cryostat sections were collected at 40X in a confocal microscope. Right panels show enlargements of boxed regions in left panels. Arrowheads in (A,C, right panels) indicate the same four cells. E-G) Immunofluorescence for calbindin (E-F) and a control with the primary antibody omitted (G-H) on 50 nm sections of amygdala. E, G) 10X images of deplasticized sections. F,H) 60X images of deplasticized sections (left), the same region on an adjacent section labeled with the plastic intact (center), and LR White sections (right). Arrows in (F) indicate the same cell in adjacent deplasticized and plastic sections. LA: lateral amygdala; Ba: basal amygdala; Ce: central amygdala; St: striatum; CB: calbindin; PV: parvalbumin. Scale bar = 100 µm in (A-D, left panels, E) and 20 µm in (A-D, right panels, F, H).

**Figure S4.**
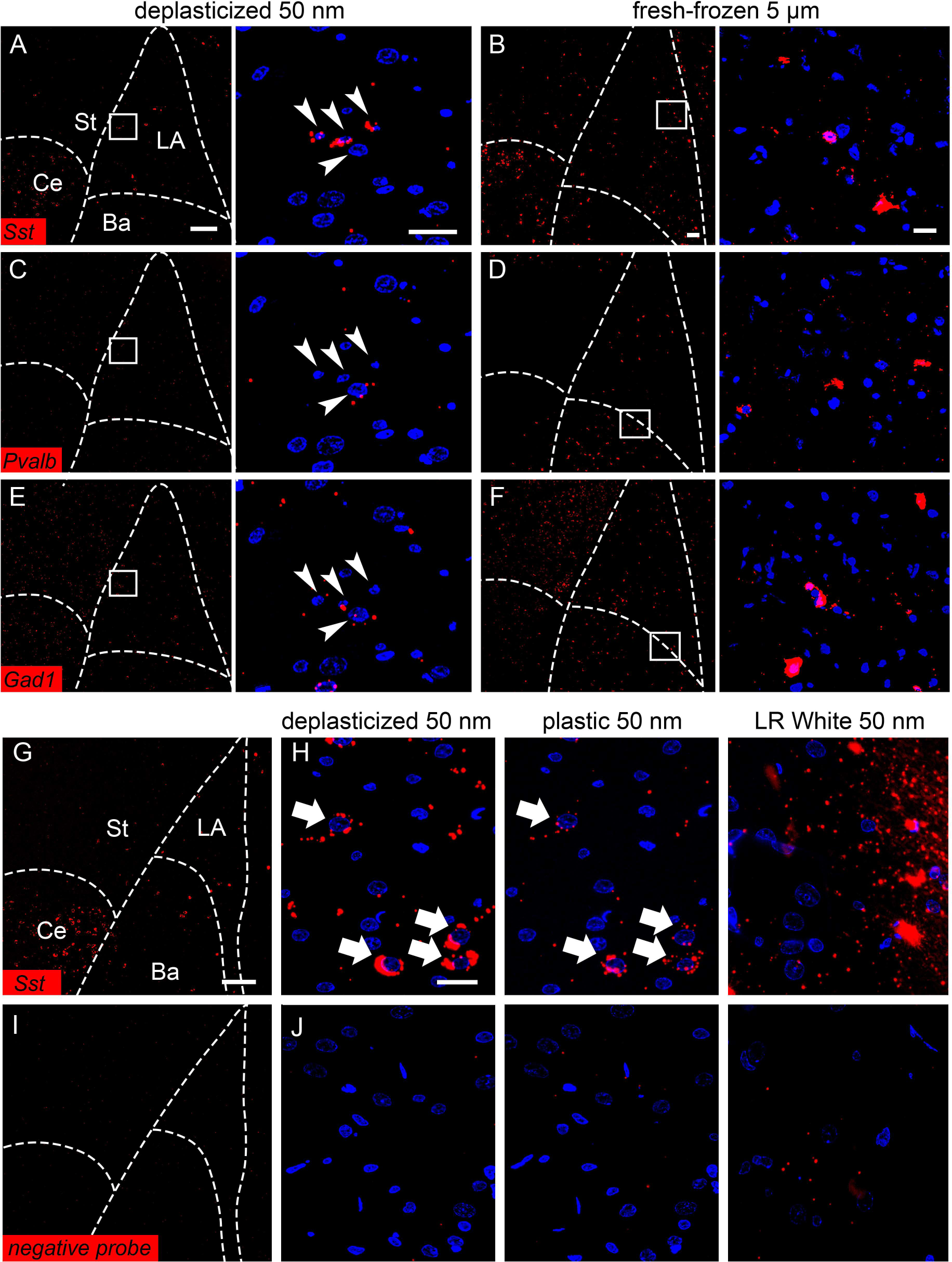
FISH labeling on deplasticized sections and standard cryosections. A-F) FISH labeling with probes for *Sst* (A,B), *Pvalb* (C,D), and *Gad1* (E,F) on deplasticized 50 nm sections (A,C,E) and 5 µm cryosections of fresh-frozen tissue (B,D,F). Ultrathin sections were imaged at 20X on a widefield microscope and single 0.5 µm optical sections of cryostat sections were collected at 40X in a confocal microscope. Right panels show enlargements of boxed regions in left panels. Arrowheads in (A,C,E, right panels) indicate the same four cells. G-J) FISH labeling with an *Sst* probe (G-H) and a negative control probe (I-J) on 50 nm sections. G,I) 10X images of deplasticized sections. H,J) 60X images of deplasticized sections (left), the same region on an adjacent section labeled with the plastic intact (center), and LR White sections (right). Arrows in (H) indicate the same cell in adjacent deplasticized and plastic sections. LA: lateral amygdala; Ba: basal amygdala; Ce: central amygdala; St: striatum. Scale bar = 100 µm in (A-F, left panels, G,I) and 20 µm in (A-F, right panels, H, J).

**Figure S5.**
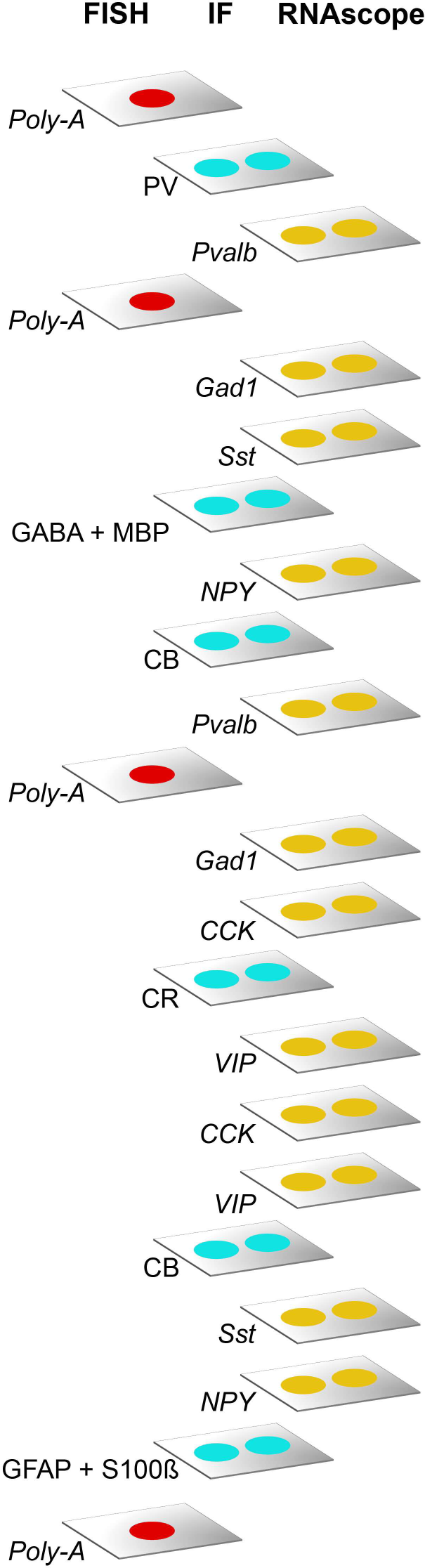
Ultraplex microscopy labeling workflow. For multiplex labeling using antibodies and RNA probes, individual 40 serial ultrathin 50 nm sections were placed on glass coverslips as shown. Sections on coverslips are processed with specific labeling protocols and reagents for each marker used.

**Figure S6.**
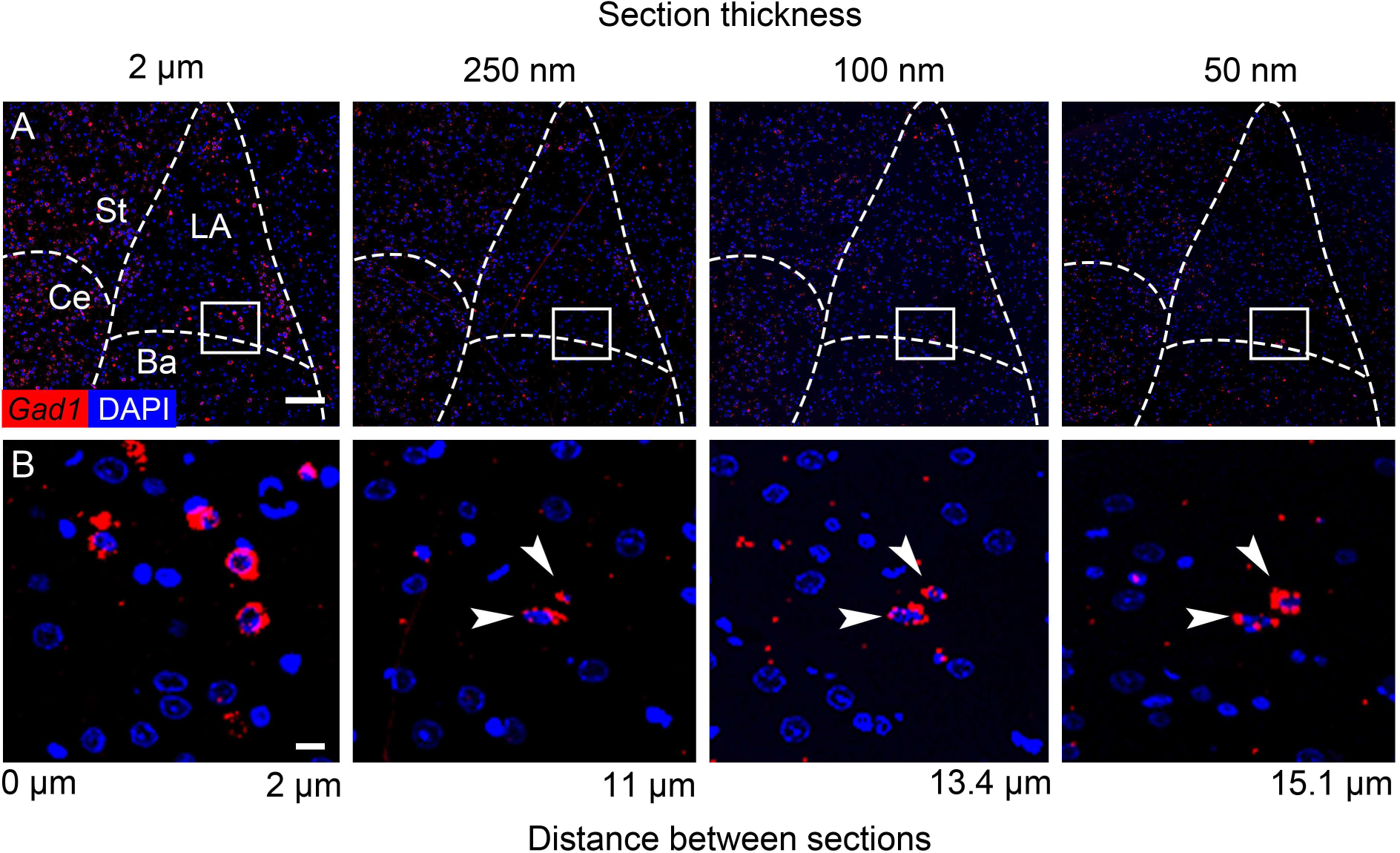
FISH labeling on deplasticized sections of different thicknesses. Non-consecutive sections spanning a total depth of 15.1 µm were labeled with a probe for *Gad1* mRNA and imaged at 20X. The lower panels show the boxed region in the top panels. Labeling density within individual cells was higher on 2 µm sections than on thinner sections, and labeled cells (arrowheads) visible on the 50 – 250 nm sections had similar labeling density across thicknesses. LA: lateral amygdala; Ba: basal amygdala; Ce: central amygdala; St: striatum; *Gad1*: glutamate decarboxylase 1 mRNA. Scale bar = 100 µm in (A), and 10 µm in (B).

**Figure S7.**
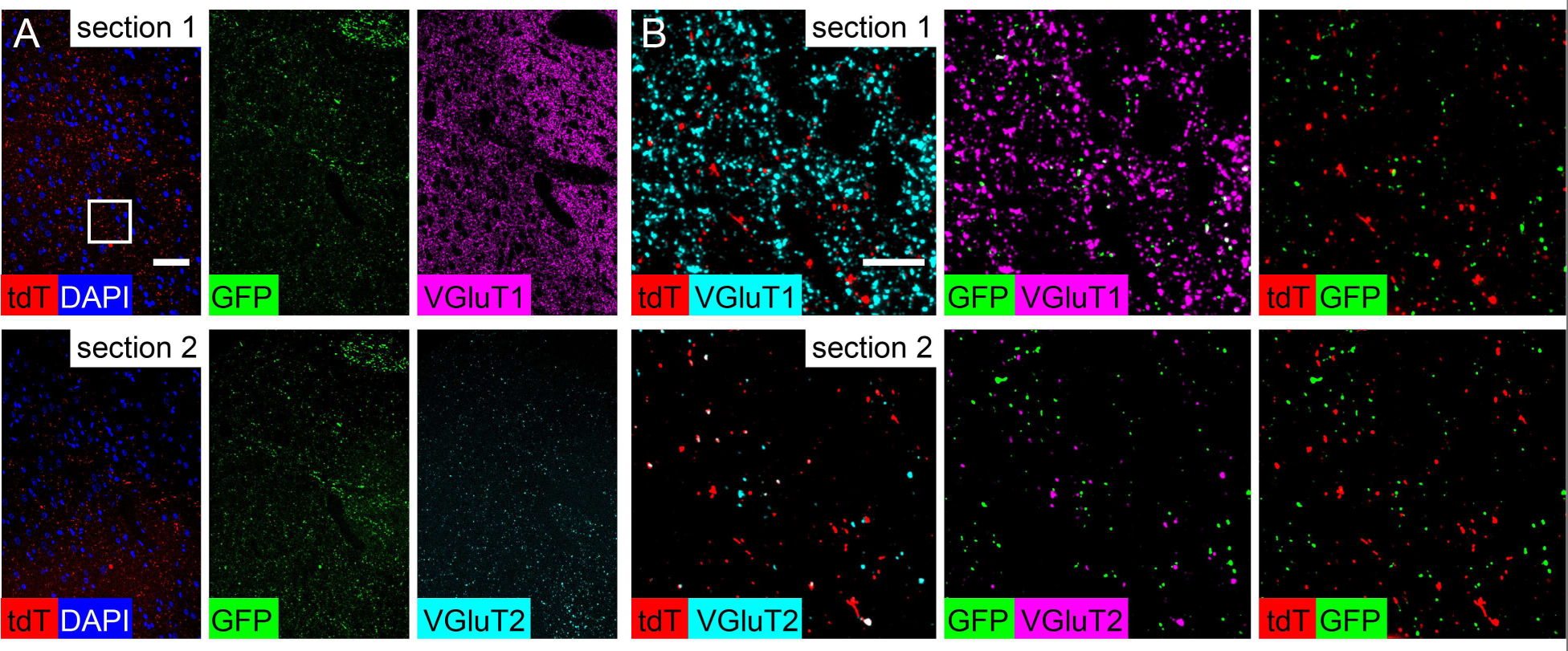
High-resolution co-localization of native fluorescence with immunofluorescence in axons. Two adjacent 50 nm sections of amygdala with thalamic axons expressing tdTomato (tdT) and cortical axons expressing GFP, one immunolabeled for VGluT1 (section 1, upper panels) and one for VGluT2 (section 2, lower panels). The same field in both sections is shown at 20X in (A), and the boxed region is shown at 100X in (B). Scale bar = 50 µm in (A, left panels), 10 µm in (B, left panels).

## References

1. Yuste, R., Hawrylycz, M., Aalling, N., Aguilar-Valles, A., Arendt, D., Arnedillo, R.A., Ascoli, G.A., Bielza, C., Bokharaie, V., Bergmann, T.B., et al. (2020). A community-based transcriptomics classification and nomenclature of neocortical cell types. Nat Neurosci 23, 1456–1468. 10.1038/s41593-020-0685-8.

2. Peng, H., Xie, P., Liu, L., Kuang, X., Wang, Y., Qu, L., Gong, H., Jiang, S., Li, A., Ruan, Z., et al. (2021). Morphological diversity of single neurons in molecularly defined cell types. Nature 598, 174–181. 10.1038/S41586-021-03941-1.

3. Sun, C., and Schuman, E.M. (2022). Logistics of neuronal protein turnover: Numbers and mechanisms. Mol Cell Neurosci 123. 10.1016/J.MCN.2022.103793.

4. Bolognesi, M.M., Manzoni, M., Scalia, C.R., Zannella, S., Bosisio, F.M., Faretta, M., and Cattoretti, G. (2017). Multiplex Staining by Sequential Immunostaining and Antibody Removal on Routine Tissue Sections. Journal of Histochemistry and Cytochemistry 65, 431–444. 10.1369/0022155417719419.

5. Klevanski, M., Herrmannsdoerfer, F., Sass, S., Venkataramani, V., Heilemann, M., and Kuner, T. (2020). Automated highly multiplexed super-resolution imaging of protein nano-architecture in cells and tissues. Nat Commun 11. 10.1038/s41467-020-15362-1.

6. Lee, J., Yoo, M., and Choi, J. (2022). Recent advances in spatially resolved transcriptomics: challenges and opportunities. BMB Rep 55, 113–124. 10.5483/BMBREP.2022.55.3.014.

7. Lin, J.R., Fallahi-Sichani, M., and Sorger, P.K. (2015). Highly multiplexed imaging of single cells using a high-throughput cyclic immunofluorescence method. Nat Commun 6. 10.1038/ncomms9390.

8. Schubert, W., Bonnekoh, B., Pommer, A.J., Philipsen, L., Böckelmann, R., Malykh, Y., Gollnick, H., Friedenberger, M., Bode, M., and Dress, A.W.M. (2006). Analyzing proteome topology and function by automated multidimensional fluorescence microscopy. Nat Biotechnol 24, 1270–1278. 10.1038/nbt1250.

9. Saka, S.K., Wang, Y., Kishi, J.Y., Zhu, A., Zeng, Y., Xie, W., Kirli, K., Yapp, C., Cicconet, M., Beliveau, B.J., et al. (2019). Immuno-SABER enables highly multiplexed and amplified protein imaging in tissues. Nat Biotechnol 37, 1080–1090. 10.1038/s41587-019-0207-y.

10. McMahon, N.P., Jones, J.A., Kwon, S., Chin, K., Nederlof, M.A., Gray, J.W., and Gibbs, S.L. (2020). Oligonucleotide conjugated antibodies permit highly multiplexed immunofluorescence for future use in clinical histopathology. J Biomed Opt 25, 1. 10.1117/1.jbo.25.5.056004.

11. Nehmé, B., Henry, M., and Mouginot, D. (2011). Combined fluorescent in situ hybridization and immunofluorescence: Limiting factors and a substitution strategy for slide-mounted tissue sections. J Neurosci Methods 196, 281–288. 10.1016/j.jneumeth.2011.01.018.

12. Moritz, C.P., Mühlhaus, T., Tenzer, S., Schulenborg, T., and Friauf, E. (2019). Poor transcript-protein correlation in the brain: negatively correlating gene products reveal neuronal polarity as a potential cause. J Neurochem 149, 582–604. 10.1111/jnc.14664.

13. Bauernfeind, A.L., and Babbitt, C.C. (2017). The predictive nature of transcript expression levels on protein expression in adult human brain. BMC Genomics 18, 322. 10.1186/s12864-017-3674-x.

14. Willingham, M.C. (1983). An alternative fixation-processing method for preembedding ultrastructural immunocytochemistry of cytoplasmic antigens: The GBS (glutaraldehyde-borohydride-saponin) procedure. Journal of Histochemistry and Cytochemistry 31, 791–798. 10.1177/31.6.6404984.

15. Sesack, S., Miner, L., and Omelchenko, N. (2006). Preembedding immunoelectron microscopy: applications for studies of the nervous system. In Neuroanatomical Tract-Tracing 3, L. Zaborszky, F. Wouterlood, and L. J. Luis, eds. (Springer), pp. 6–71.

16. Kosaka, T., Kosaka, K., Tateishi, K., Hamaoka, Y., Yanaihara, N., Wu, J.-Y., and Hama, K. (1985). GABAergic neurons containing CCK-8-like and/or VIP-like immunoreactivities in the rat hippocampus and dentate gyrus. J Comp Neurol 239, 420–430. 10.1002/cne.902390408.

17. McDonald, A.J., and Mascagni, F. (2001). Colocalization of calcium-binding proteins and GABA in neurons of the rat basolateral amygdala. Neuroscience 105, 681–693.

18. Talapka, P., Kocsis, Z., Marsi, L.D., Szarvas, V.E., and Kisvárday, Z.F. (2021). Application of the Mirror Technique for Three-Dimensional Electron Microscopy of Neurochemically Identified GABA-ergic Dendrites. Front Neuroanat 15, 21. 10.3389/fnana.2021.652422.

19. Harris, K.M., Perry, E., Bourne, J., Feinberg, M., Ostroff, L., and Hurlburt, J. (2006). Uniform serial sectioning for transmission electron microscopy. Journal of Neuroscience 26, 12101–12103. 10.1523/JNEUROSCI.3994-06.2006.

20. Zheng, Z., Lauritzen, J.S., Perlman, E., Robinson, C.G., Nichols, M., Milkie, D., Torrens, O., Price, J., Fisher, C.B., Sharifi, N., et al. (2018). A complete electron microscopy volume of the brain of adult Drosophila melanogaster. Cell 174, 730–743.e22. 10.1016/j.cell.2018.06.019.

21. Kasthuri, N., Hayworth, K.J., Berger, D.R., Schalek, R.L., Conchello, J.A., Knowles-Barley, S., Lee, D., Vázquez-Reina, A., Kaynig, V., Jones, T.R., et al. (2015). Saturated reconstruction of a volume of neocortex. Cell 162, 648–661. 10.1016/j.cell.2015.06.054.

22. Pfeiffer, R.L., Anderson, J.R., Dahal, J., Garcia, J.C., Yang, J.H., Sigulinsky, C.L., Rapp, K., Emrich, D.P., Watt, C.B., Johnstun, H.A., et al. (2020). A pathoconnectome of early neurodegeneration: Network changes in retinal degeneration. Exp Eye Res 199. 10.1016/J.EXER.2020.108196.

23. Ottersen, O.P., Storm-Mathisen, J., and Somogyi, P. (1988). Colocalization of glycine-like and GABA-like immunoreactivities in Golgi cell terminals in the rat cerebellum: a postembedding light and electron microscopic study. Brain Res 450, 342–353. 10.1016/0006-8993(88)91573-9.

24. Sato, F., Nakamura, Y., and Shinoda, Y. (1997). Serial electron microscopic reconstruction of axon terminals on physiologically identified thalamocortical neurons in the cat ventral lateral nucleus. Journal of Comparative Neurology 388, 613–631. 10.1002/(SICI)1096-9861(19971201)388:4<613::AID-CNE9>3.0.CO;2-5.

25. Shahidi, R., Williams, E.A., Conzelmann, M., Asadulina, A., Verasztó, C., Jasek, S., Bezares-Calderón, L.A., and Jékely, G. (2015). A serial multiplex immunogold labeling method for identifying peptidergic neurons in connectomes. Elife 4, e11147. 10.7554/eLife.11147.

26. Merighi, A., Polak, J.M., Fumagalli, G., and Theodosis, D.T. (1989). Ultrastructural localization of neuropeptides and GABA in rat dorsal horn: A comparison of different immunogold labeling techniques. Journal of Histochemistry and Cytochemistry 37, 529–540. 10.1177/37.4.2564404.

27. Brorson, S.H., and Skjørten, F. (1995). Mechanism for antigen detection on deplasticized epoxy sections. Micron 26, 301–310. 10.1016/0968-4328(95)00008-9.

28. Litwin, J.A. (1985). Light Microscopic Histochemistry on Plastic Sections. Prog Histochem Cytochem 16, 3–5. 10.1016/S0079-6336(85)80001-2.

29. Grube, D., and Kusumoto, Y. (1986). Serial Semithin Sections in Immunohistochemistry: Techniques and applications. Archivum histologicum japonicum 49, 391–410. 10.1679/aohc.49.391.

30. Petralia, R.S., and Wang, Y.X. (2021). Review of Post-embedding Immunogold Methods for the Study of Neuronal Structures. Front Neuroanat 15. 10.3389/fnana.2021.763427.

31. Newman, G.R., and Hobot, J.A. (1987). Modern acrylics for post-embedding immunostaining techniques. Journal of Histochemistry and Cytochemistry 35, 971–981. 10.1177/35.9.3302021.

32. Holderith, N., Heredi, J., Kis, V., and Nusser, Z. (2020). A High-Resolution Method for Quantitative Molecular Analysis of Functionally Characterized Individual Synapses. Cell Rep 32. 10.1016/j.celrep.2020.107968.

33. Micheva, K.D., and Smith, S.J. (2007). Array tomography: a new tool for imaging the molecular architecture and ultrastructure of neural circuits. Neuron 55, 25–36. 10.1016/j.neuron.2007.06.014.

34. Avila, A.S., and Henstridge, C.M. (2022). Array tomography: 15 years of synaptic analysis. Neuronal Signal 6. 10.1042/NS20220013.

35. De Camilli, P., Cameron, R., and Greengard, P. (1983). Synapsin I (protein I), a nerve terminal-specific phosphoprotein. I. Its general distribution in synapses of the central and peripheral nervous system demonstrated by immunofluorescence in frozen and plastic sections. Journal of Cell Biology 96, 1337– 1354. 10.1083/jcb.96.5.1337.

36. Micheva, K.D., Gong, B., Collman, F., Weinberg, R.J., Smith, S.J., Trimmer, J.S., and Murray, K.D. (2023). Developing a Toolbox of Antibodies Validated for Array Tomography-Based Imaging of Brain Synapses. eNeuro 10, ENEURO.0290-23.2023. 10.1523/ENEURO.0290-23.2023.

37. Mansfield, J.R. (2017). Phenotyping multiple subsets of immune cells in situ in FFPE tissue sections: An overview of methodologies. In Methods in Molecular Biology (Humana Press Inc.), pp. 75–99. 10.1007/978-1-4939-6730-8_5.

38. Fuller, C.E., and Perry, A. (2002). Fluorescence in situ hybridization (FISH) in diagnostic and investigative neuropathology. Brain Pathol 12, 67–86. 10.1111/J.1750-3639.2002.TB00424.X.

39. Newman, G.R., and Hobot, J.A. (1999). Resins for combined light and electron microscopy: A half century of development. Histochemical Journal 31, 495–505. 10.1023/A:1003850921869.

40. Holmes, P.F., Bohrer, M., and Kohn, J. (2008). Exploration of polymethacrylate structure-property correlations: Advances towards combinatorial and high-throughput methods for biomaterials discovery. Prog Polym Sci 33, 787. 10.1016/J.PROGPOLYMSCI.2008.05.002.

41. Parfitt, G.J. (2019). Immunofluorescence Tomography: High-resolution 3-D reconstruction by serial-sectioning of methacrylate embedded tissues and alignment of 2-D immunofluorescence images. Sci Rep 9, 1–9. 10.1038/s41598-018-38232-9.

42. Saito, C., Hayashi, M., Sakai, A., Fujie, M., Kuroiwa, H., and Kuroiwa, T. (1999). Improved sensitivity for high resolution in situ hybridization using resin extraction of methyl methacrylate embedded material. Biotechnic and Histochemistry 74, 40–48. 10.3109/10520299909066476.

43. Warren, K.C., Coyne, K.J., Waite, J.H., and Cary, S.C. (1998). Use of methacrylate de-embedding protocols for in situ hybridization on semithin plastic sections with multiple detection strategies. Journal of Histochemistry and Cytochemistry 46, 149–155. 10.1177/002215549804600203.

44. Torgersen, J.S., Takle, H., and Andersen, Ø. (2009). Localization of mRNAs and proteins in methyl methacrylate-embedded tissues. Journal of Histochemistry and Cytochemistry 57, 825–830. 10.1369/jhc.2009.953695.

45. McDonald, A.J. (1989). Coexistence of somatostatin with neuropeptide Y, but not with cholecystokinin or vasoactive intestinal peptide, in neurons of the rat amygdala. Brain Res 500, 37–45.

46. McDonald, A.J. (2020). Functional neuroanatomy of the basolateral amygdala: Neurons, neurotransmitters, and circuits. In Handbook of Behavioral Neuroscience (Elsevier B.V.), pp. 1–38. 10.1016/B978-0-12-815134-1.00001-5.

47. Wang, F., Flanagan, J., Su, N., Wang, L.C., Bui, S., Nielson, A., Wu, X., Vo, H.T., Ma, X.J., and Luo, Y. (2012). RNAscope: a novel in situ RNA analysis platform for formalin-fixed, paraffin-embedded tissues. J Mol Diagn 14, 22–29. 10.1016/J.JMOLDX.2011.08.002.

48. Zhou, H., Gang, Y., Chen, S., Wang, Y., Xiong, Y., Li, L., Yin, F., Liu, Y., Liu, X., and Zeng, S. (2017). Development of a neutral embedding resin for optical imaging of fluorescently labeled biological tissue. J Biomed Opt 22, 1. 10.1117/1.JBO.22.10.106015.

49. Watanabe, S., Punge, A., Hollopeter, G., Willig, K.I., Hobson, R.J., Davis, M.W., Hell, S.W., and Jorgensen, E.M. (2011). Protein localization in electron micrographs using fluorescence nanoscopy. Nat Methods 8, 80–84. 10.1038/nmeth.1537.

50. Torrealba, F., and Carrasco, M.A. (2004). A review on electron microscopy and neurotransmitter systems. Brain Res Rev 47, 5–17. 10.1016/j.brainresrev.2004.06.004.

51. Roberts, G.W., Woodhams, P.L., Polak, J.M., and Crow, T.J. (1982). Distribution of neuropeptides in the limbic system of the rat: The amygdaloid complex. Neuroscience 7, 99–131. 10.1016/0306-4522(82)90156-7.

52. Marley, P.D., Emson, P.C., Hunt, S.P., and Fahrenkrug, J. (1981). A long ascending projection in the rat brain containing vasoactive intestinal polypeptide. Neurosci Lett 27, 261–266. 10.1016/0304-3940(81)90440-7.

53. Cassell, M.D., Gray, T.S., and Williams, T.H. (1982). Changes in the distribution of peptidergic terminals in the central nucleus of the rat amygdala following lesions of its medial input. Anatomical Record 202, 28A.

54. Gray, T.S., Cassell, M.D., Nilaver, G., Zimmerman, E.A., and Williams, T.H. (1984). The distribution and ultrastructure of VIP-immunoreactivity in the central nucleus of the rat amygdala. Neuroscience 11, 399–408. 10.1016/0306-4522(84)90032-0.

55. Micheva, K.D., O’Rourke, N., Busse, B., and Smith, S.J. (2010). Array tomography: immunostaining and antibody elution. Cold Spring Harb Protoc 2010, pdb.prot5525.

56. Newman, G.R. (1987). Use and abuse of LR White. Histochem J 19, 118–120. 10.1007/BF01682756.

57. Kramer, S., Meyer-Natus, E., Stigloher, C., Thoma, H., Schnaufer, A., and Engstler, M. (2020). Parallel monitoring of RNA abundance, localization and compactness with correlative single molecule FISH on LR White embedded samples. Nucleic Acids Res 49, 14. 10.1093/nar/gkaa1142.

58. Matsuno, A., Kirino, T., Ohsugi, Y., Utsunomiya, H., Takekoshi, S., Osamura, R.Y., Watanabe, K., and Teramoto, A. (1994). Ultrastructural distribution of growth hormone and prolactin mRNAs in normal rat pituitary cells: a comparison between preembedding and postembedding methods. Histochemistry 102, 265–270. 10.1007/BF00269162.

59. Ukimura, A., Deguchi, H., Kitaura, Y., Fujioka, S., Hirasawa, M., Kawamura, K., and Hirai, K. (1997). Intracellular viral localization in murine Coxsackievirus-B3 myocarditis: Ultrastructural study by electron microscopic in situ hybridization. American Journal of Pathology 150, 2061–2074.

60. Ostroff, L.E.L.E., Santini, E., Sears, R., Deane, Z., Kanadia, R.N.R.N., Ledoux, J.E.J.E., Lhakhang, T., Tsirigos, A., Heguy, A., and Klann, E. (2019). Axon TRAP reveals learning-associated alterations in cortical axonal mRNAs in the lateral amgydala. Elife 8. 10.7554/eLife.51607.

61. Souquere, S., Mollet, S., Kress, M., Dautry, F., Pierron, G., and Weil, D. (2009). Unravelling the ultrastructure of stress granules and associated P-bodies in human cells. J Cell Sci 122, 3619–3626. 10.1242/jcs.054437.

62. Dörries, U., Bartsch, U., Nolte, C., Roth, J., and Schachner, M. (1993). Adaptation of a non-radioactive in situ hybridization method to electron microscopy: detection of tenascin mRNAs in mouse cerebellum with digoxigenin-labelled probes and gold-labelled antibodies. Histochemistry 99, 251–262. 10.1007/BF00269143.

63. Troxler, M., Egger, D., Pfister, T., and Bienz, K. (1992). Intracellular localization of poliovirus RNA by in Situ hybridization at the ultrastructural level using single-stranded riboprobes. Virology 191, 687–697. 10.1016/0042-6822(92)90244-J.

64. Dirks, R.W., Van Dorp, A.G.M., Van Minnen, J., Fransen, J.A.M., Van der Ploeg, M., and Raap, A.K. (1992). Electron microscopic detection of RNA sequences by non-radioactive in situ hybridization in the mollusk Lymnaea stagnalis. Journal of Histochemistry and Cytochemistry 40, 1647–1657. 10.1177/40.11.1431053.

65. Griffiths, Gareth. (1993). Fine Structure Immunocytochemistry (Springer Berlin Heidelberg).

66. Erickson, P.A., Anderson, D.H., and Fisher, S.K. (1987). Use of uranyl acetate en bloc to improve tissue preservation and labeling for post-embedding immunoelectron microscopy. J Electron Microsc Tech 5, 303–314. 10.1002/jemt.1060050403.

67. McDonald, K. (1999). High-pressure freezing for preservation of high resolution fine structure and antigenicity for immunolabeling. Methods Mol Biol 117, 77–97. 10.1385/1-59259-201-5:77.

68. Quintana, C. (1994). Cryofixation, cryosubstitution, cryoembedding for ultrastructural, immunocytochemical and microanalytical studies. Micron 25, 63–99.

69. van Lookeren Campagne, M., Oestreicher, A.B., van der Krift, T.P., Gispen, W.H., and Verkleij, A.J. (1991). Freeze-substitution and Lowicryl HM20 embedding of fixed rat brain: suitability for immunogold ultrastructural localization of neural antigens. Journal of Histochemistry & Cytochemistry 39, 1267–1279. 10.1177/39.9.1833448.

70. Armbruster, B.L., Carlemalm, E., Chiovetti, R., Garavito, R.M., Hobot, J.A., Kellenberger, E., and Villiger, W. (1982). Specimen preparation for electron microscopy using low temperature embedding resins. J Microsc 126, 77–85. 10.1111/j.1365-2818.1982.tb00358.x.

71. Micheva, K.D., O’Rourke, N., Busse, B., and Smith, S.J. (2010). Array tomography: production of arrays. Cold Spring Harb Protoc 2010, pdb.prot5524.

72. Zhou, H., Xiong, Y., Wang, Y., Wang, X., Li, P., Gang, Y., Liu, X., and Zeng, S. (2017). High-refractive index of acrylate embedding resin clarifies mouse brain tissue. J Biomed Opt 22, 1. 10.1117/1.JBO.22.11.110503.

73. Delage, E., Guilbert, T., and Yates, F. (2023). Reproducibility: Successful 3D imaging of cleared biological samples with light sheet fluorescence microscopy. J Cell Biol 222. 10.1083/JCB.202307143.

74. Schindelin, J., Arganda-Carreras, I., Frise, E., Kaynig, V., Longair, M., Pietzsch, T., Preibisch, S., Rueden, C., Saalfeld, S., Schmid, B., et al. (2012). Fiji: an open-source platform for biological-image analysis. Nat Methods 9, 676–682. 10.1038/nmeth.2019.

75. Manders, E., Verbeek, F., and Aten, J. (1993). Measurement of colocalization of objects in dual-color confocal images. J Microsc Oxford 169, 375–382.

