## Supplementary material for "Ultraplex microscopy: versatile highly-multiplexed molecular labeling and imaging across scale and resolution": Key Resources Table

| Reagent or Resource | Source | Identifier | Additional information |
|---|---|---|---|
| **Antibodies** | | | |
| anti-Calbindin (CB) (rabbit polyclonal) | Synaptic Systems | 214 003, lot 1-3, RRID:AB_2619901 | 1:1000 (plastic/deplasticized), 1:500 (free floating) |
| anti-Calcitonin Gene-Related Peptide (CGRP) (guinea pig polyclonal) | Synaptic Systems | 414 004, lot 1-5, RRID:AB_2737049 | 1: 5000 (deplasticized) |
| anti-Calretinin (CR) (rabbit polyclonal) | Synaptic Systems | 214 102, lot 1-5, RRID:AB_2228331 | 1:1000 (deplasticized) |
| anti-DsRed (Rabbit polyclonal) | Takarabio | 632496, lot 2103116, RRID:AB_10013483 | 1:1000 (deplasticized) |
| anti-Glial Fibrillary Acidic Protein (GFAP) (mouse monoclonal) | Sigma Aldrich | G3893, lot 0000182427, RRID:AB_477010 | 1:1000 (deplasticized) |
| anti-Green Fluorescent Protein (GFP) (Chicken polyclonal) | Abcam | ab13970, lot GR236651-25, RRID:AB_300798 | 1:1000 (deplasticized) |
| anti-Microtubule-associated protein 2 (map2) (chicken polyclonal) | Abcam | ab5392, lot 1049576-2, RRID:AB_2138153 | 1:1000 (deplasticized) |
| anti-Myelin Basic Protein (MBP) (Mouse monoclonal) | Biolegend | 808401, lot B315597, RRID:AB_2564742 | 1:1000 (plastic/deplasticized) |
| anti-Neurofilament (SMI312) (Mouse monoclonal) | Biolegend | 837904, lot B333596, RRID:AB_2566782 | 1:1000 (deplasticized) |
| anti-Parvalbumin (PV) (rabbit polyclonal) | Abcam | ab11427, lot 1029943-5, RRID: AB_298032 | 1:1000 (plastic/deplasticized), 1:500 (free floating) |
| anti-S100 Beta (S100B) (rabbit polyclonall) | Proteintech | 15146-1-AP, lot 00103243, RRID:AB_2254244 | 1:1000 (deplasticized) |
| anti-somatostatin (SOM) (chicken polyclonal) | Synaptic Systems | 366 006, lot 1-7, RRID:AB_2636910 | 1:500 (deplasticized) |
| anti-Vasoactive Intentinal Peptide (VIP) (guinea pig polyclonal) | Synaptic Systems | 443 005, lot 1-5, RRID:AB_2832228 | 1:1000 (deplasticized), 1:100 (free floating) |
| anti-Vesicular Glutamate Transporter 1 (vGluT1) (rabbit polyclonal) | Synaptic Systems | 135 303, lot 4-107, RRID:AB_887875 | 1:1000 (deplasticized) |
| anti-Vesicular Glutamate Transporter 2 (vGluT2) (guinea pig polyclonal) | Synaptic Systems | 135 404, lot 3-47 RRID:AB_887884 | 1:10,000 (deplasticized) |
| anti-γ-Aminobutyric acid (GABA) (rabbit polyclonal) | Sigma Aldrich | A2052, lot, 0000177418, RRID:AB_477652 | 1:10,000 (deplasticized) |
| Lectin fro Wisteria floribunda (WFA) (biotin) | Sigma Aldrich | L1516, lot SLCM8716 | 1:1000 (deplasticized) |
| Goat anti-Chicken Alexa Fluor 488 (Secondary) | Invitrogen | A32931, lot XB343360, RRID:AB_2762843 | 1:200 (deplasticized) |

| | | | |
|---|---|---|---|
| Goat anti-Chicken Alexa Fluor 647 (Secondary) | Invitrogen | A21449, lot 2480082, RRID:AB_2535866 | 1:200 (deplasticized) |
| Goat anti-Guinea pig Alexa Fluor 488 (Secondary) | Invitrogen | A11073, lot 2160428, RRID:AB_2534117 | 1:200 (deplasticized) |
| Goat anti-Guinea pig Alexa Fluor 647 (Secondary) | Invitrogen | A21450, lot 2446026, RRID:AB_2535867 | 1:200 (deplasticized) |
| Goat anti-Mouse Alexa Fluor 488 (Secondary) | Invitrogen | A11001, lot 2659299, RRID:AB_2534069 | 1:200 (deplasticized) |
| Goat anti-Mouse Alexa Fluor 647 (Secondary) | Invitrogen | A21235, lot 2482945, RRID:AB_2535804 | 1:200 (deplasticized) |
| Goat anti-Rabbit Alexa Fluor 568 (Secondary) | Invitrogen | A11011, lot 2379475, RRID:AB_143157 | 1:200 (deplasticized) |
| Goat anti-Rabbit Alexa Fluor 647 (Secondary) | Invitrogen | A21245, lot 2641997, RRID:AB_2535813 | 1:200 (deplasticized/free floating) |
| Goat anti-Rabbit Alexa Fluor Plus 488 (Secondary) | Invitrogen | A32731, lot XD343356, RRID:AB_2633280 | 1:200 (deplasticized/free floating) |
| STAR Orange Rabbit | Abberior | STORANGE-1002 | 1:200 (deplasticized) |
| STAR Red Mouse | Abberior | STRED-1001 | 1:200 (deplasticized) |
| Streptavidin, Alexa Fluor 488 (Secondary) | Invitrogen | S32354, lot 2387462 | 1:200 (deplasticized) |
| Streptavidin, Alexa Fluor 647 (Secondary) | Invitrogen | S32357, lot 2581824 | 1:200 (deplasticized) |
| **Sequence-based reagent** | | | |
| *Cck* ISH probe | Advanced Cell Diagnostics | 532851-C1 | Multiplexed FISH |
| *DapB* ISH probe (negative) | Advanced Cell Diagnostics | 310043-C1 | Multiplexed FISH |
| *Gad1* ISH probe | Advanced Cell Diagnostics | 316401-C1 | Multiplexed FISH |
| *Npy-O1* ISH probe | Advanced Cell Diagnostics | 1211911-C1 | Multiplexed FISH |
| *Pvalb* ISH probe | Advanced Cell Diagnostics | 407821-C1 | Multiplexed FISH |
| *Sst* ISH probe | Advanced Cell Diagnostics | 412181-C1 | Multiplexed FISH |
| *Vip* ISH probe | Advanced Cell Diagnostics | 485681-C1 | Multiplexed FISH |

| | | | |
|---|---|---|---|
| *Poly-A* ISH probe | Integrated DNA Technologies | Ostroff et al., 2019 | 2 µM (deplasticized) |
| **Recombinant DNA reagent** | | | |
| pENN.AAV1.hSyn.Cre.WPRE.hGH | Addgene | 105553-AAV1 | N/A |
| pAAV8-FLEX-tdTomato | Addgene | 28306-AAV8 | N/A |
| pAAV8-hSyn-EGFP | Addgene | 50465-AAV8 | N/A |
| pAAV8-hSyn-mCherry | Addgene | 114472-AAV8 | N/A |
| **Chemicals or Reagents** | | | |
| RNAscope H2O2 and Protease reagents | Advanced Cell Diagnostics | 322381 | Multiplexed FISH |
| RNAscope Multiplex Fluorescent Detection Reagents V2 | Advanced Cell Diagnostics | 323110 | Multiplexed FISH |
| RNAscope Wash Buffer reagents | Advanced Cell Diagnostics | 310091 | Multiplexed FISH |
| RNAscope Multiplex TSA Buffer | Advanced Cell Diagnostics | 322810 | Multiplexed FISH |
| 4′,6-Diamidine-2′-phenylindole dihydrochloride (DAPI) | Millipore Sigma | 10236276001 | 2 µg/ml |
| TSA Vivid 650 | Advanced Cell Diagnostics | 323273 | 1:500 - Multiplexed FISH |
| Xyazine 100mg/mL | Covertus | NDC 1165-4024-1 | Surgical Sedative |
| Zetamine (Ketamine Hydrochloride) 100mg/mL | VET ONE | NDC 13985-702-10 | Surgical Sedative |
| Chloral hydreate (C-IVN) | Sigma Aldrich | C8383-250G | Perfusion setative, mixed into 25% solution in water |
| Ketofen (ketoprofen) 100mg/mL | Zoetis | NDC 54771-4396-1 | Surgical and post-surgical non-steroidal anti-inflammatory |
| Paraformaldehyde, granular | Electron Microscopy Sciences | 19210 | N/A |
| Glutaraldehyde 50% aqueous | Electron Microscopy Sciences | 16320 | N/A |
| Sodium Phosphate, Dibasic Heptahydrate | Electron Microscopy Sciences | 21182 | N/A |

| | | | |
|---|---|---|---|
| Sodium Phosphate, Monobasic | Electron Microscopy Sciences | 21190 | N/A |
| Sodium Citrate | Electron Microscopy Sciences | 21140 | N/A |
| Sodium Chloride, granular | Fisher Scientific | S6403 | N/A |
| Uranyl Acetate | Structure Probe | 02624-AB | N/A |
| Polyvinylpyrrolidone | Millipore Sigma | 5295 | N/A |
| LR White | Electron Microscopy Sciences | 14383-UC | N/A |
| Methyl Methacrylate, Monomer | Electron Microscopy Sciences | 18800 | N/A |
| n-Butyl Methacrylate, Monomer | Electron Microscopy Sciences | 12100 | N/A |
| Benzoin Methyl Ether, UV Catalyst | Electron Microscopy Sciences | 11290 | N/A |
| Triton X-100 | Electron Microscopy Sciences | 22140 | N/A |
| Tween-20 | Sigma Aldrich | P1379 | N/A |
| Bovine Serum Albumin | Jackson Immunoresearch | 001-000-162 | |
| Hematoxylin and Eosin Stain Kit | Vector Laboratories | H-3502 | N/A |
| Sodium Tetraborate | Sigma Aldrich | 221732 | N/A |
| Toluidine Blue | Electron Microscopy Sciences | 22050 | N/A |
| Acetone, Reagent Grade | Electron Microscopy Sciences | 10014 | N/A |
| Ethyl Alcohol, Anhydrous 200 proof | Electron Microscopy Sciences | 15056 | N/A |
| Ultrapure Dnase/Rnase Free Distilled Water | Thermo Scientific | 10977023 | N/A |
| Pioloform B Resin | SPI-CHEM | 63148-65 | N/A |
| **Other** | | | |

| | | | |
|---|---|---|---|
| Synaptek NOTCH Grids | Electron Microscopy Sciences | S2010-NOTCH | N/A |
| Flat Embedding Capsules, Polyethylene | Electron Microscopy Sciences | 70021 | N/A |
| Gelatin Capsules | Electron Microscopy Sciences | 70103 | N/A |
| ProLong Gold Antifade Mountant | Invitrogen | P36930 | N/A |
| Gelatin, Type A, 300 Bloom | Electron Microscopy Sciences | 16564 | N/A |
| Chromium (III) potassium sulfate dodecahydrate | Sigma Aldrich | 243361 | N/A |
| Superfrost Disposable Microscope Slides | Fisher Scientific | 12-550-143 | N/A |
| Cover-GOLD SEAL Coverslips | Electron Microscopy Sciences | 63792 | N/A |
| Superfrost Plus Gold Slides | Fisher Scientific | 15-188-48 | N/A |
| µ-Slide 1 Well Glass Bottom coverslip | Ibidi | 82107 | N/A |
| Pap Pen ImmEdge Hydrophobic Barrier Pen | Vector Laboratories | H-4000 | N/A |
| HybEZTM II oven | Advanced Cell Diagnostics | PN 321710 | N/A |
| Humidity Control Tray | Advanced Cell Diagnostics | PN 310012 | N/A |
| Perfect loop | Electron Microscopy Sciences | 70939 | N/A |
| LEGATO® 101 SYRINGE PUMP | KD Scientific | 78-8101 | Syringe pump cannula injections |
| Vibratome VT1200 S | Leica | N/A | N/A |
| Ultramicrotome EM UCT | Leica | N/A | N/A |
| Cryostat, CM 3050 S | Leica | N/A | N/A |
| AFS2 Freeze Substitution System | Leica | N/A | N/A |
| Model 962 Dual Ultra Precise Small Animal Stereotaxic Instrument | Kopf | N/A | Stereotaxic Mount |

| | | | |
|---|---|---|---|
| Ni-E Eclipse Nikon Motorized Upright Epifluorecence Microscope | Nikon | N/A | N/A |
| Nikon Ti2E Inverted Microscopes | Nikon | N/A | N/A |
| Abberior STED Inverted Olympus IX83 Microscope | Abberior Instruments | N/A | N/A |
| Leica SP8 Spectral Confocal | Leica | N/A | N/A |
| Leica Thunder Imager | Leica | N/A | N/A |
| JEOL 1400 Transmission Electron Micrsocope | JEOL | N/A | N/A |
| **Software** | | | |
| ImageJ, Version 2.14.0/1.54f | National Intitute of Health | RRID: SCR_003070 | N/A |
| Adobe Photoshop, Version 25.3.1 | Adobe | N/A | N/A |
| Statistica, Version 13 | Tibco | | |
